# An improved nanobody-based approach to capture and visualize specific interaction networks of binary protein complexes in living cells

**DOI:** 10.1101/2024.09.12.612471

**Authors:** Nawal Hajj Sleiman, Julie Carnesecchi, Yunlong Jia, Fréderic Delolme, Laurent Gilquin, Patrice Gouet, Samir Merabet

**Affiliations:** Institut de Génomique Fonctionnelle de Lyon, CNRS UMR5242, ENS de Lyon, Université Lyon I, Lyon, France; Institut de Génétique Moléculaire de Montpellier, CNRS, Université de Montpellier, Montpellier, France; Department of Developmental and Cellular Biology, University of California Irvine, Irvine, CA 92697, USA; Protein Science Facility, CNRS-UAR3444, INSERM-US8, École Normale Supérieure de Lyon, Université Lyon I, Lyon, France; Molecular Microbiology and Structural Biochemistry, CNRS UMR5086, Université Lyon I, Lyon, France

**Keywords:** Bi-nano-ID, BiFC, GFP-nanobody, TurboID, TAZ, Hippo, SERPINB3/4

## Abstract

Protein interaction networks (or interactomes) are formed progressively, each interaction influencing the next one. Accordingly, a same protein will establish different interactomes depending on its first associated cofactor, thereby diversifying its function in the cell. In contrast to their central role, few methods exist to capture interactomes of dimeric protein complexes. Here, we tackle this issue by introducing an innovative method based on bimolecular fluorescence complementation and the specific binding of a nanobody fused to a proximity-dependent biotinylating enzyme. This method was applied to visualize and capture specific interactions of the cytoplasmic TAZ/14-3-3e and nuclear TAZ/TEAD2 complexes, which are major downstream effectors of the Hippo signaling pathway. Among other interactions, we identified SERPINB4 as a novel regulator of TAZ and 14-3-3e proliferative activity in mesenchymal stromal cells. Molecular dissections in living cells revealed the central role of a unique residue of TAZ for recruiting SERPINB4 specifically in the presence of 14-3-3e. Overall, our work demonstrates the importance of considering binary protein complexes for deciphering interactomes and establishes a novel sensitive method in this perspective.

## Introduction

Proteins do not work individually within the cell environment, but by forming connected networks, or interactomes, with other protein-binding partners. These molecular networks are established by successive protein-protein interactions (PPIs) and will vary depending on the cell type and/or cell compartment. This molecular versatility is at the basis of protein function diversification [1]. In addition, transient and low-affinity PPIs are central for shaping interactome identity *in vivo*. Therefore, capturing weak PPIs has become a technological challenge for understanding protein function in normal or pathological conditions [2].

The conventional technique to identify endogenous interactomes of a protein of interest (POI) is affinity purification (AP) and the identification of interacting proteins by liquid chromatography followed by tandem-mass spectrometry (LC/MS-MS). However, AP-based approaches are biased towards direct, high affinity and stable PPIs [3]. The recent development of several proximity-dependent labeling (PL) methods in living cells partially addressed the difficulty to capture low affinity and transient PPIs. These methods take advantage of promiscuous enzymes fused to the POI that can label proximal endogenous proteins (within a radius of approximately 10 nanometers), including transient and weak interaction partners. Although different types of labeling reactions have been described, the most widespread approaches rely on the covalent binding of the small biotin molecule, allowing single-step streptavidin AP before subsequent MS analysis of labelled proteins [3]. One of the recent and most commonly used biotin-based proximity labeling enzymes is the TurboID, which is a directed-evolution variant of the original BioID, with less toxicity, smaller size and considerably faster labeling activity [4]. Split forms of TurboID (split-TurboID) have also been developed to capture interactions of a dimeric protein complex, allowing probing the proteomic composition in a subcellular compartment [5] or cell surface environment [6,7] and identifying SUMO-dependent interactions [8]. However, the activity of the reconstituted TurboID is relatively low compared to the full length/uncut form [5]. Consequently, it requires long incubation times with the biotin (24 hours), which forbids having access to interactome temporal dynamics. Moreover, biotin ligases do not allow observing PPIs in a living cell context.

In this work, we develop a methodology called Bi-nano-ID (**bi**molecular fluorescent complementation (BiFC)-directed **nano**body coupled to proximity **id**entification enzyme). This method aims at specifically characterizing endogenous interactomes of a specific dimeric bait protein complex. Bi-nano-ID was also established with a combination of fluorescent protein fragments that are compatible for doing bicolor BiFC. This property allows to simultaneously visualize the interaction of the novel candidate partner together with the dimeric bait protein complex in the living cell.

As a proof of concept, Bi-nano-ID was established by using TAZ (Transcriptional Coactivator with PDZ-binding motif, also named WWTR1) associated to two different cofactors. TAZ (and another functionally related protein called Yes-associated protein, YAP) plays pivotal roles as a downstream component of the Hippo signaling pathway, with a well-known nucleocytoplasmic shuttling property in cells [9]. When the Hippo signaling is active, upstream kinases phosphorylate TAZ, leading to its retention in the cytoplasm and interaction with 14-3-3 proteins. Cytoplasmic TAZ/14-3-3 complexes can promote proliferation and the differentiation of various tissue types [10]. Upon Hippo signaling inactivation, TAZ translocates into the nucleus and interacts with TEAD proteins to bind DNA, resulting in the expression of several target genes. TAZ/TEAD complexes are required for the differentiation of osteoblasts and other cell types [11,12]. Aiming to verify the applicability of Bi-nano-ID, we mapped the interactome of TAZ associated with either the long isoform of 14-3-3 (14-3-3e, also named YWHAE) or the isoform 2 of TEAD (TEAD2) in human HEK293T cells. These two isoforms have been described to regulate the hippo pathway in different cell contexts, including cancers (see for example [13–15]). The role of a novel specific candidate partner of TAZ/14-3-3e complexes was further confirmed at the functional and molecular levels, revealing an unexpected level of interaction plasticity in TAZ and 14-3-3e proteins.

Overall, our work demonstrates that Bi-nano-ID is a sensitive method for deciphering interactomes of specific binary protein complexes in living cells.

## Results

### Capturing the generic interactome of TAZ in living HEK293T cells

Given that we aimed at establishing Bi-nano-ID by considering TAZ associated with either a cytoplasmic or a nuclear cofactor in HEK293T cells, we first asked for the general interaction properties of TAZ in the same cell context. To this end, we generated a TAZ-TurboID fusion construct under the control of a doxycycline (dox)-inducible promoter (see Material and Methods). We assumed that proteins that will be biotinylated by TAZ-TurboID will represent the generic interactome of TAZ, therefore all possible interactions without considering their dependency towards a specific TAZ-associated cofactor (Fig. 1A). In addition, getting the global interactome of TAZ-TurboID expressed in HEK293T cells was also a prerequisite to confirm that this cell line was appropriate for revealing functionally relevant interactions, *i.e.* interactions involved in the Hippo signaling pathway.

**Figure 1:**
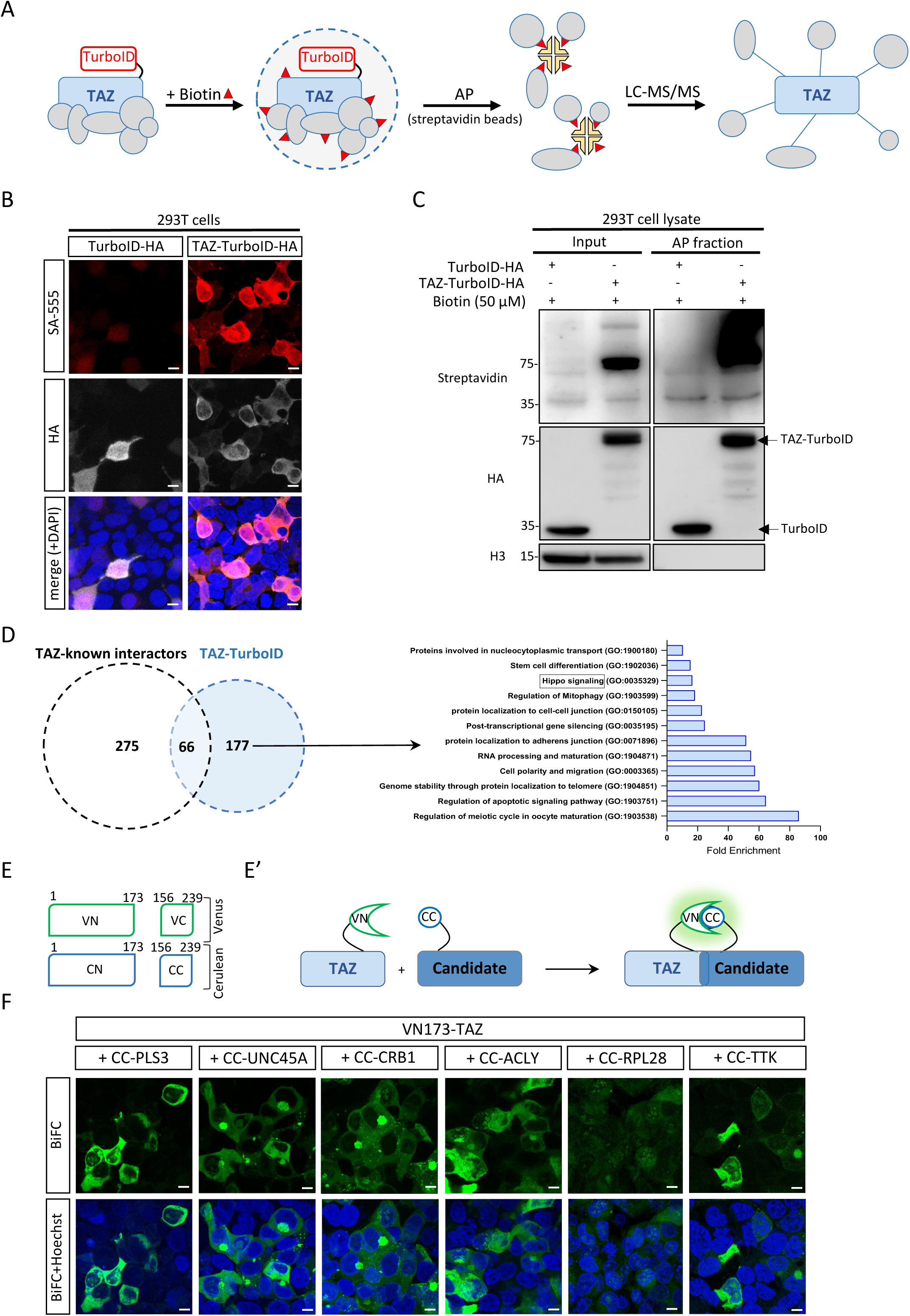
Capturing the generic interactome of TAZ-TurboID in HEK-293T cells. **(A)** Schematic representation of the capture of TAZ-TurboID interactome. The addition of Biotin leads to the labeling of proteins in proximity to the bait protein construct. Biotinylated proteins are affinity purified with streptavidin-coupled beads and analyzed by liquid chromatography coupled to tandem mass spectrometry (LC-MS/MS). **(B)** Validation of the specific biotinylating activity of TAZ-TurboID when compared to TurboID alone by immunofluorescence in HEK293T cells. Biotinylation was revealed with streptavidin (SA) coupled to the 555 dyes (red). The expression of each construct was confirmed with an immunostaining against the HA tag present in each construct (gray). Dapi (blue) stains for nuclei. Scale bar = 20 µm. **(C)** Western blot analysis of biotin labeling by TurboID or TAZ-TurboID. The input and affinity purified (AP) biotinylated proteins from samples expressing each construct were analyzed via western blotting after biotin incubation (10 minutes). Samples were probed by streptavidin conjugate, HA and Histone 3 (H3) antibodies. The band corresponding to TurboID or TAZ-TurboID is indicated with anti-HA. **(D)** GO term analysis of the TAZ-TurboID interactome compared to the known interactions of TAZ in the literature. The 177 novel interactions captured in this study have several GO term enrichments directly (box) or indirectly related to the Hippo signaling pathway. **(E-E’)** Principle of BiFC with Venus and Cerulean derived fragments. The N-terminal fragment of Venus (VN) can complement with the C-terminal fragment of Cerulean (CC) and provide a Venus-like fluorescence signal. **(F)** Illustrative confocal acquisitions of BiFC between VN-Taz and new candidate partners fused to CC, as indicated. Hoechst stains for nuclei in live HEK293T cells (blue). Scale bar = 20 µm.

The expression and activity of TAZ-TurboID was first verified by fluorescent immunostaining (Fig. 1B). Control of specificity was performed with the expression of TurboID alone. This experiment showed strong biotinylation activity in the context of the fusion with TAZ, while weak signals were observed with TurboID alone (Fig.1B and S1). Staining also revealed a preferential localization and biotinylation activity of TAZ-TurboID in the cytoplasm of HEK293T cells (Fig. 1B). Western blots (WB) confirmed the specificity of TAZ-TurboID when compared to TurboID upon AP of biotinylated proteins (Fig. 1C and S1). Altogether, immunostaining and AP experiments validated the specific activity of TAZ-TurboID, and we next performed cell protein extracts for LC/MS-MS analysis with three biological replicates of TAZ-TurboID and TurboID (see also Materials and Methods).

Principal component analysis (PCA) showed a distinct distribution for TAZ-TurboID and TurboID replicates (Fig. S1). Moreover, the overlap between the three TAZ-TurboID replicates was high (between 71% to 79% of interactions were common between two replicates, and between 30% to 40% of interactions were common to the three replicates (Fig. S1).

All TAZ-TurboID interactions common to at least two different biological replicates and not found with TurboID alone were considered, leading to a total number of 243 proteins (Table S1). We compared this list to the 379 known physical interactors of TAZ described in the literature (https://thebiogrid.org/117434/summary/homo-sapiens/wwtr1.html). Most of these known physical interactors of TAZ have been identified by systematic AP-MS profiling of C-terminally FLAG-HA-tagged bait proteins expressed in HEK293T cells [16]. 66 interactions were present in the two lists. Importantly, one of the most enriched GO terms for the 66 common interactors relates to the Hippo signaling pathway (Fig. S1 and Table S1). Interestingly, biological functions associated with the hippo signaling pathway were also enriched when considering the remaining 177 interactions of TAZ-TurboID but not when considering the remaining 275 known interactions (Fig. 1D, S1 and Table S1). Other GO terms related to the hippo pathway were also enriched in the 66 common and 177 TAZ-TurboID interactions (such as cell polarity, cell-cell junction and RNA processing: Fig. 1D and S1), further reinforcing the functional relevance of the two lists.

We next selected 6 known and 6 novel TAZ interactors for performing individual Bimolecular Fluorescence Complementation (BiFC) in live HEK293T cells (see Materials and Methods). BiFC relies on the property of monomeric fluorescent proteins to be reconstituted upon complementation when the two sub-fragments are closed enough in space ([17]). It allows assessing whether two candidate proteins could be part of the same protein complex with a simple readout (apparition of a fluorescent signal upon excitation, Fig. 1E-E’). BiFC confirmed that all tested candidates could interact with TAZ. Interaction profiles were diverse although a majority were located in the cytoplasm, which was somehow consistent with the preferential localization and biotinylation activity of TAZ-TurboID in HEK293T cells (Fig. 1F and S1).

Altogether, these results showed that AP of biotinylated proteins in HEK293T cells was appropriate for revealing a generic TAZ interactome with functionally relevant GO term enrichments. The validation of the TurboID efficiency in HEK293T to capture TAZ generic interactome allowed the implementation of this biotinylation-based approach for establishing the Bi-nano-ID tool. Moreover, TAZ generic interactome will serve as a reference for further analyses of the specific interactomes of TAZ associated with its cofactors (TAZ/14-3-3e and TAZ/TEAD2 interactome).

### Establishing Bi-nano-ID experimental parameters in HEK293T cells

Bi-nano-ID is based on an anti-GFP nanobody that will specifically recognize a BiFC complex on the one side, and the coupling to the TurboID enzyme on the other side (Fig. 2A). More particularly, the nanobody (named <u>c</u>onditionally <u>s</u>table (cs) <u>G</u>FP-<u>B</u>inding <u>P</u>eptide, csGBP) has been engineered to be unstable when unbound to its epitope, allowing getting a better signal-to-noise ratio, and was originally described in the context of BiFC resulting from the complementation between the N-terminal (0-155: VN155) and C-terminal (156-239: VC) fragment of Venus (a derivative of the yellow fluorescent protein YFP, [18]). The same GFP lacking the conditionally stable mutations, has also previously been used to capture interactions of Venus-based BiFC complexes by immunoprecipitation [19,20].

**Figure 2:**
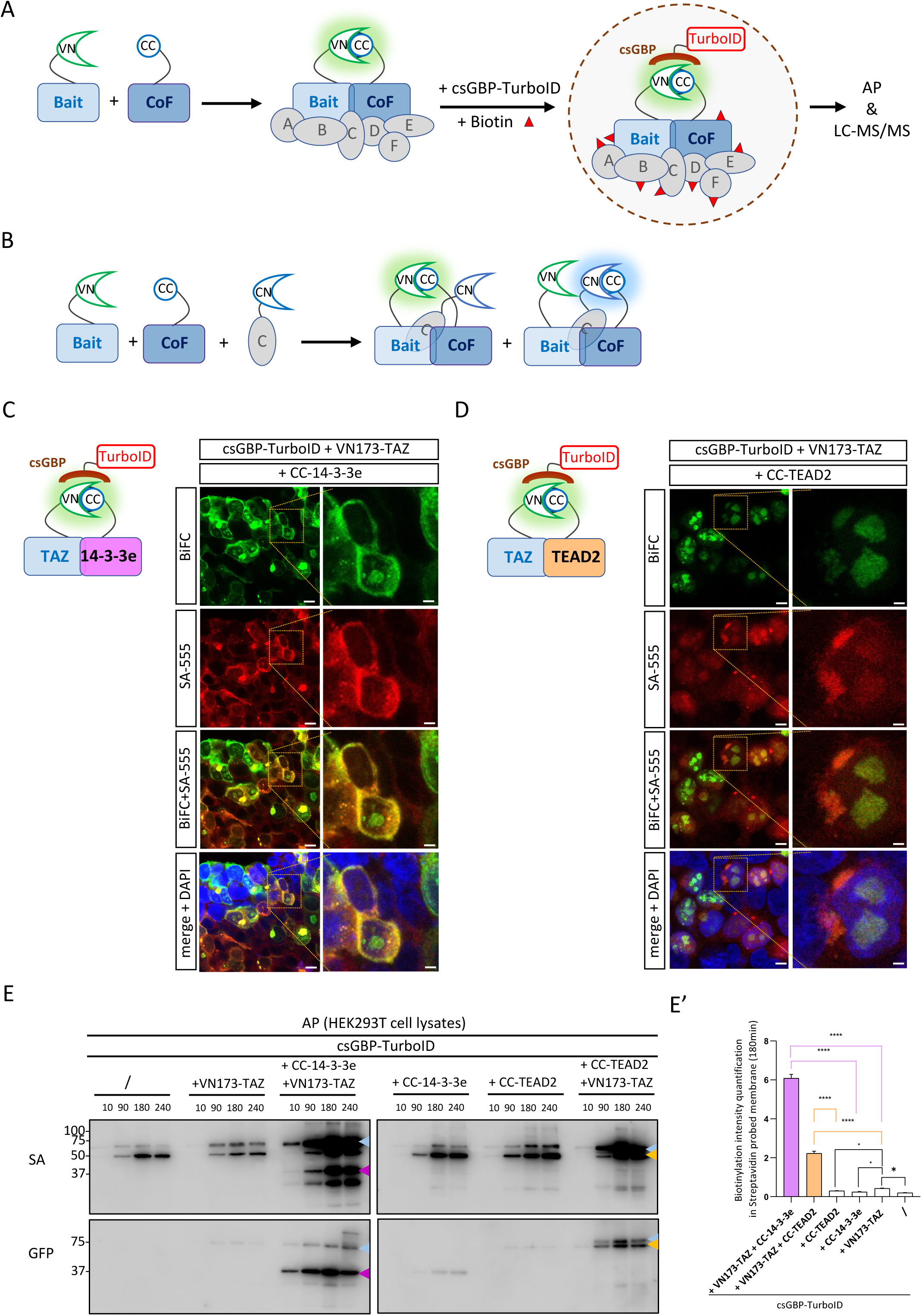
Establishing Bi-nano-ID experimental parameters in HEK293T cells. **(A)** Principle of Bi-nano-ID. BiFC resulting from the complementation between the VN and CC fragment is recognized by a GFP nanobody (csGBP) coupled to the TurboID enzyme. The addition of biotin allows the biotinylation of surrounding partners, which can then be purified by affinity purification (AP) and analyzed by mass spectrometry (LC-MS/MS). **(B)** Simultaneous visualization of the interaction between a novel candidate partner (gray ball) and the bait dimeric complex. The superposition of Venus-(green) and Cerulean-like (blue) fluorescent signals in the cell is indicative of co-localized interactions, therefore of possible trimeric complex assembly. **(C-D)** Validation of the specificity of Bi-nano-ID by immunofluorescence. Biotinylation is revealed with streptavidin (SA) coupled to 555 dyes (red). Interaction between the fusion TAZ and 14-3-3e (C) or TEAD2 (D) proteins is revealed by BiFC (green). Dapi stains for nuclei (blue). Enlargements highlight the specific colocalization of red and green signals in the cytoplasm or the nucleus for TAZ/14-3-3e and TAZ/TEAD2 complexes, respectively. Scale bar = 20 µm. **(E)** Western blot of affinity purified biotinylated proteins from samples expressing dsGBP-TurboID individually or in combination with VN173-TAZ, and/or CC-14-3-3e or CC/TEAD2. Experiment was performed after different biotin (50 µM) incubation timepoints (10, 90, 180, 240 minutes), as indicated. Samples were probes by streptavidin conjugate (to reveal biotinylated proteins) and GFP (to reveal the VN and CC fusion constructs). Colored arrowheads depict sizes corresponding to VN-TAZ (blue arrowhead), CC-14-3-3e (magenta arrowhead) and CC-TEAD2 (orange arrowhead). Note that non-specific bands with the nanobody alone are present at the size of VN-TAZ and CC-TEAD2. These bands were taken into account for the quantification of the biotinylation activity of the nanobody with VN-TAZ (upper band) and CC-TEAD2 (lower band). **(E’)** Quantification of the intensity of the biotinylation bands in the different conditions at 180min biotin incubation time, as indicated. Quantification was obtained from three biological replicates. Statistical tests were performed with one-way ANOVA (\**p* < 0.05, \*\*\*\**p* < 0.0001).

In this work, we aimed at testing the efficiency and specificity of the csGBP nanobody in the context of a complementation system that allows performing bicolor BiFC. This complementation system has been developed to visualize two different PPIs simultaneously, and could serve for the subsequent validation of novel interactions of the dimeric bait protein complex in living cells (Fig. 2B). Bicolor BiFC relies on the property of the N-terminal fragment of YFP (yellow fluorescent protein) to complement with the C-terminal fragment of the blue fluorescent protein ECFP (enhanced cyan fluorescent protein), producing a yellow-like fluorescent signal [21,22]. Two different split sites in the N-terminal fragment of YFP (at the position 155 or 173) can complement with the C-terminal fragment of ECFP (156-239), with the 1-173 N-terminal fragment described to produce stronger fluorescent signals than the 1-155 N-terminal fragment [21]. Here, we tested the two complementation sites by using the Venus and Cerulean fluorescent proteins, which display faster maturation time and higher fluorescence brightness than YFP and ECFP, respectively (Fig. S2 and [23,24]). Practically, TAZ was fused to the 155-or 173 N-terminal fragment of Venus (making VN155-TAZ and VN173-TAZ constructs), and the 14-3-3e and TEAD2 cofactors were fused to the 156-239 C-terminal fragment of Cerulean (making CC-14-3-3e and CC-TEAD2 constructs). All constructs were under the same dox-inducible promoter and BiFC was analyzed in living HEK293T cells. Results showed that BiFC obtained with the VN155 fragment was extremely weak when compared to BiFC obtained with VN173, underlining that VN155 could not complement efficiently with the CC fragment in both TAZ/14-3-3e and TAZ/TEAD2 complexes (Fig. S2). BiFC with VN173-TAZ also validated the correct cytoplasmic or nuclear localization of TAZ/14-3-3e and TAZ/TEAD2 complexes in live HEK293T cells, respectively (Fig. S2). Collectively, these experiments confirmed that the VN155 fragment was not best appropriate for doing bicolor BiFC and we thus decided to establish Bi-nano-ID by using the VN173 fragment.

The specificity of the csGBP-TurboID nanobody was first evaluated by immunofluorescence. BiFC constructs were co-transfected with csGBP-TurboID encoding plasmid, induced with dox, and biotin was added before visualizing both BiFC and biotinylation (see Materials and Methods). These analyses confirmed that BiFC and biotinylation signals colocalized in the case of TAZ/14-3-3e and TAZ/TEAD2 complexes (Fig. 2C-D). In contrast, no biotinylation could be observed when expressing the csGBP-TurboID nanobody alone or together with the individual VN173-TAZ, CC-14-3-3e or CC-TEAD2 construct, demonstrating that the biotinylating activity was dependent on the binding of the nanobody to BiFC complexes (Fig. S3). Biotinylation signals were also extremely weak in the case of BiFC complexes with VN155-TAZ, underlining that the binding affinity of the nanobody was dependent on the complementation efficiency between the N-and C-terminal fragments (Fig. S2).

Next, we tested the specificity of our tools upon affinity purification (AP) with streptavidin-coupled beads. We applied the same BiFC and control conditions as described for the immunostaining. Proteins were extracted, purified and revealed after different incubation timepoints with the biotin to select the best experimental condition for having strong BiFC-specific biotinylation signals with a good signal-to-noise ratio. All quantifications were performed by normalizing the AP over the input and the correct expression of each construct was also validated by using an anti-GFP recognizing the VN and CC fragments (Fig.S4). Results showed non-specific biotinylation activity of the nanobody in the absence of any BiFC constructs (first four lanes in Fig. 2E). Importantly, signals were strongly enriched in the presence of BiFC complexes and quantifications showed that the best incubation time was 180 min, with no stronger BiFC-specific biotinylation signals at 240 min (Fig. 2E-E’ and S4). The specificity of the nanobody was also more precisely quantified by considering each single construct. This analysis showed negligeable biotinylation activity in the case of CC-14-3-3e and CC-TEAD2 constructs (in comparison to csGBP-TurboID alone: Fig. 2E-E’ and S4). In contrast, more signal was observed with VN-TAZ alone (in comparison to the csGBP-TurboID alone: Fig. 2E-E’ and S4), indicating that the csGBP-TurboID nanobody was weakly recognizing the VN173 fragment. In conclusion, these experiments established the experimental conditions for purifying BiFC-specific interactomes with the csGBP-TurboID nanobody tool. Because of the weak recognition of the VN173 fragment by the nanobody, we also considered VN-TAZ as an additional filtering control for the analysis of BiFC-specific MS data (see below).

### Using Bi-nano-ID to capture specific interactomes of TAZ/14-3-3e and TAZ/TEAD2 complexes in living HEK293T cells

Mass spectrometry analysis was performed by using the streptavidin affinity purified fraction of protein extracts from HEK293T cells in the different control and BiFC conditions. Control conditions consisted in the expression of the nanobody alone (to discriminate for the general background of the TurboID enzyme, referred to as csGBP-TurboID), and to the co-expression of csGBP-TurboID with VN-TAZ (to discriminate for the weak affinity of the nanobody to the VN173 fragment). These two control conditions were used to filter interactions obtained in the two BiFC conditions (csGBP-TurboID co-expressed with VN-TAZ and CC-14-3-3e or with VN-TAZ and CC-TEAD2, Fig. S5).

Results showed a good reproducibility between the three biological replicates in each experimental condition (between 26% to 51% of interactions were common to at least two replicates, and between 17% to 30% were common to the three replicates: Table S2 and Fig. S5). Interactions present in two or three replicates were considered for subsequent analyses. Interactions with BiFC complexes were filtered with the interactions obtained in the two control conditions, which led to the identification of 96 and 42 candidate interactions for TAZ/14-3-3e and TAZ/TEAD2 complexes, respectively (Materials and Methods, Table S2 and Fig. S5).

To better discriminate for TAZ/14-3-3e-and TAZ/TEAD2-dependent interactions, we compared the two BiFC interactomes with the interactome obtained with TAZ-TurboID. This analysis revealed 50 and 30 interactions that were specifically found with TAZ/14-3-3e and TAZ/TEAD2 BiFC complexes, respectively (Fig. 3A and S6). The finding of BiFC-specific interactions underlined that endogenous 14-3-3 and TEAD proteins did not hid all specific binary complex interactions (although half of the 96 interactions of the TAZ/14-3-3e BiFC complex were also captured with TAZ-TurboID).

**Figure 3:**
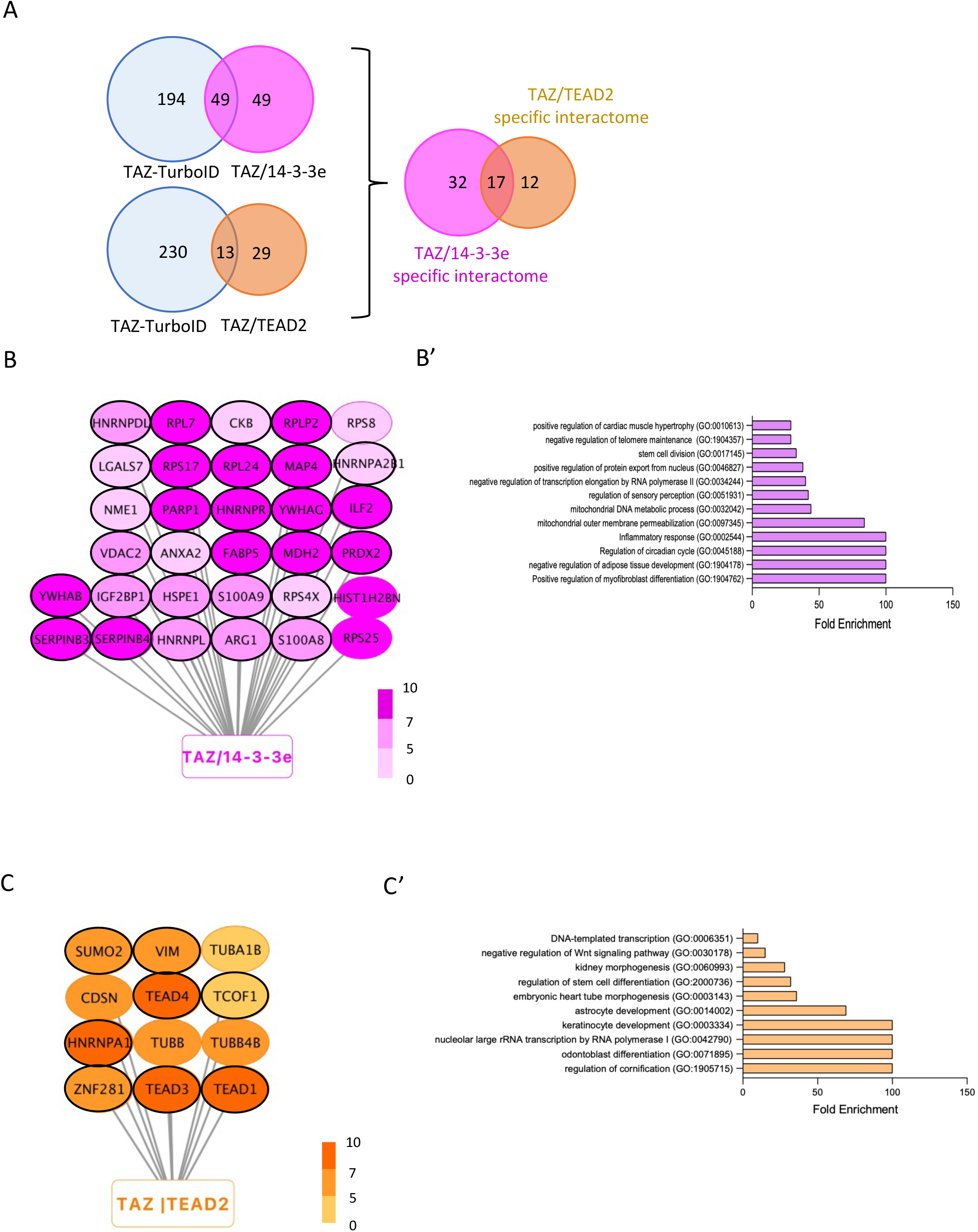
TAZ/14-3-3e and TAZ/TEAD2 interactome clustering. **(A)** Venn diagrams showing common and specific interactions between TAZ-TurboID, TAZ/14-3-3e and TAZ/TEAD2 interactomes. **(B-B’)** Specific interactions and GO term enrichment for TAZ/14-3-3e interactome. Light-to-dark pink gradient illustrates low-to-high protein enrichment, respectively (as deduced from the LFQ value of mass spectrometry data, see also Table S2). Cytoplasmic proteins are surrounded (black circle). **(C-C’)** Specific interactions and GO term enrichment for TAZ/TEAD2-interactome. Light-to-dark orange gradient illustrates low-to-high protein enrichment, respectively (as deduced from the LFQ value of mass spectrometry data, see also Table S2). Nuclear proteins are surrounded (black circle).

Importantly, the goal of Bi-nano-ID was to capture interactions that will be specifically distributed between the cytoplasmic TAZ/14-3-3e and nuclear TAZ/TEAD2 complexes. The comparison of these two interactomes revealed 32 interactions that were specific of TAZ/14-3-3e and 12 interactions that were specific of TAZ/TEAD2 (Fig. 3A-C and S6). Thus, we concluded that Bi-nano-ID was successful for capturing BiFC-specific interactomes.

In order to know the functions that could be associated with the common and/or specific TAZ-related interactomes, we performed a GO-term enrichment analysis (see Materials and Methods and Table S2). “Hippo signaling pathway” was strongly enriched in TAZ/14-3-3e and TAZ/TEAD2 interactomes (in addition to the TAZ-TurboID interactome), emphasizing that our tools led to the capture of functionally relevant interactions (Fig. 3B’-C’ and S7). Interestingly, functions related to mRNA stability and catabolic processes were also enriched in the three interactomes, suggesting that they also represent a major and generic molecular facet of TAZ (Fig. 3B’-C’ and S7). Several GO terms were specifically associated with TAZ/14-3-3e or TAZ/TEAD2 interactome. Importantly, some of these specific functions were consistent with the known activity of 14-3-3 and TEAD proteins with YAP/TAZ effectors (such as nuclear export and stem cell division for TAZ/14-3-3e, or nuclear import and transcription for TAZ/TEAD2, Fig. 3B’-C’). Other specific functions (like “negative regulation of adipose tissue development” and “circadian rhythm” for TAZ/14-3-3e, or “keratinocyte development” and “odontoblast differentiation” for TAZ/TEAD2) are associated with YAZ/TAZ in the literature (see for example [25–28]). These observations underline that Bi-nano-ID was efficient for capturing novel interactions that could be involved in known specific TAZ/14-3-3e and TAZ/TEAD2 functions. Finally, not all the generic TAZ functions were captured with the TAZ/14-3-3e and TAZ/TEAD2 interactomes (like oocyte maturation or mitophagy regulation, Fig. S7), suggesting that these TAZ functions could rely on other 14-3-3-and TEAD-independent partnerships.

### SERPINB3 and B4 potentiate the proliferative activity of TAZ and 14-3-3e in HS27A mesenchymal stromal cells

YAP/TAZ regulators have context-dependent roles in stem cells, either driving self-renewal or differentiation depending on the identity of the stem cell [29,30]. These differential activities are also linked to the cytoplasmic or nuclear localization, therefore to the association with cytoplasmic (including 14-3-3) or nuclear (including TEAD) cofactors. Here, we wondered whether some specific TAZ/14-3-3e or TAZ/TEAD2 interactions captured with Bi-nano-ID could be functionally relevant in a stem cell-like context. To this end, we considered the HS27A bone marrow mesenchymal stromal cell line that can be induced into different cell types depending on the cell culture medium [31,32]. Given that a common enriched GO term for TAZ/14-3-3e and TAZ/TEAD2 complexes was related to the “positive regulation of osteoblast differentiation” (Fig. S7), we decided to cultivate HS27A cells in an osteoblast-differentiation medium. The differentiation into osteoblast was confirmed with Alizarin red staining (see Materials and Methods and Fig. 4A-A’). We hypothesized that cofactors working in concert with TAZ/14-3-3e or TAZ/TEAD2 complexes could potentially display distinct expression profiles depending on the mesenchymal-like/undifferentiated or osteoblast-like/differentiated state of HS27A cells. In order to identify such candidate cofactors, we performed RNA-seq experiments of HS27A cells at day 1 (proliferation state) and at day 15 (osteoblast differentiation state). RNA-seq data showed several genes that were strongly up-or down-regulated when comparing the proliferative and differentiated state of HS27A cells (Fig. 4B and Table S3). Unexpectedly, neither *TAZ*, nor *TEAD* or *14-3-3* genes showed a significant differential expression level between the undifferentiated and differentiated cell state (Table S3). This observation suggests that their role could depend on post-transcriptional regulatory process, such as the regulation of the nuclear shuttling. Nevertheless, although a number of the genes displaying a differential expression profile could correspond to downstream effectors of TAZ/14-3-3e and TAZ/TEAD2 complexes, we compared the list of differentially expressed genes to our lists of specific candidate cofactors captured with Bi-nano-ID. This analysis led to the identification of *SERPINB3* and *SERPINB4.* Interestingly, these two genes were among the most downregulated genes during the differentiation process, and the corresponding proteins were specifically captured with TAZ/14-3-3e (LC/MS-MS analysis could not discriminate between SERPINB3 and B4, which share 92% identity in their amino acid sequence, Fig. 3B and S8). SERPINB3 and B4 belong to the family of serine/cysteine protease inhibitors and are frequently co-expressed in different tissues. Their physiological role remains largely unknown, but their abnormal elevated expression level has been reported to contribute to several pathologies, including chronic liver diseases, inflammatory diseases and cancer (such as squamous cell carcinomas and primary liver cancers ([33,34]). SERPINB4 has also been associated with psoriasis severity that is characterized by epidermal hyperproliferation and infiltration of inflammatory cells [35]. Moreover, the analysis of the known interactome of SERPINB4 revealed SERPINB3 and two other interaction partners that were specifically captured with TAZ/14-3-3e: S100A8 and S100A9 (Fig. 3B-C and 4C). These two proteins have been described to be part of a common circuit with the hippo signaling and promote the proliferation and colony formation of breast cancer cells [36]. Finally, “stem cell division” was also among the most enriched GO terms specific of the TAZ/14-3-3e interactome (Fig. 3B and Table S2). Altogether, these observations suggest that SERPINB3 and B4 could play a role for the proliferation of HS27A cells, making them strong candidates to test in the context of TAZ/14-3-3e proliferative activity.

**Figure 4.**
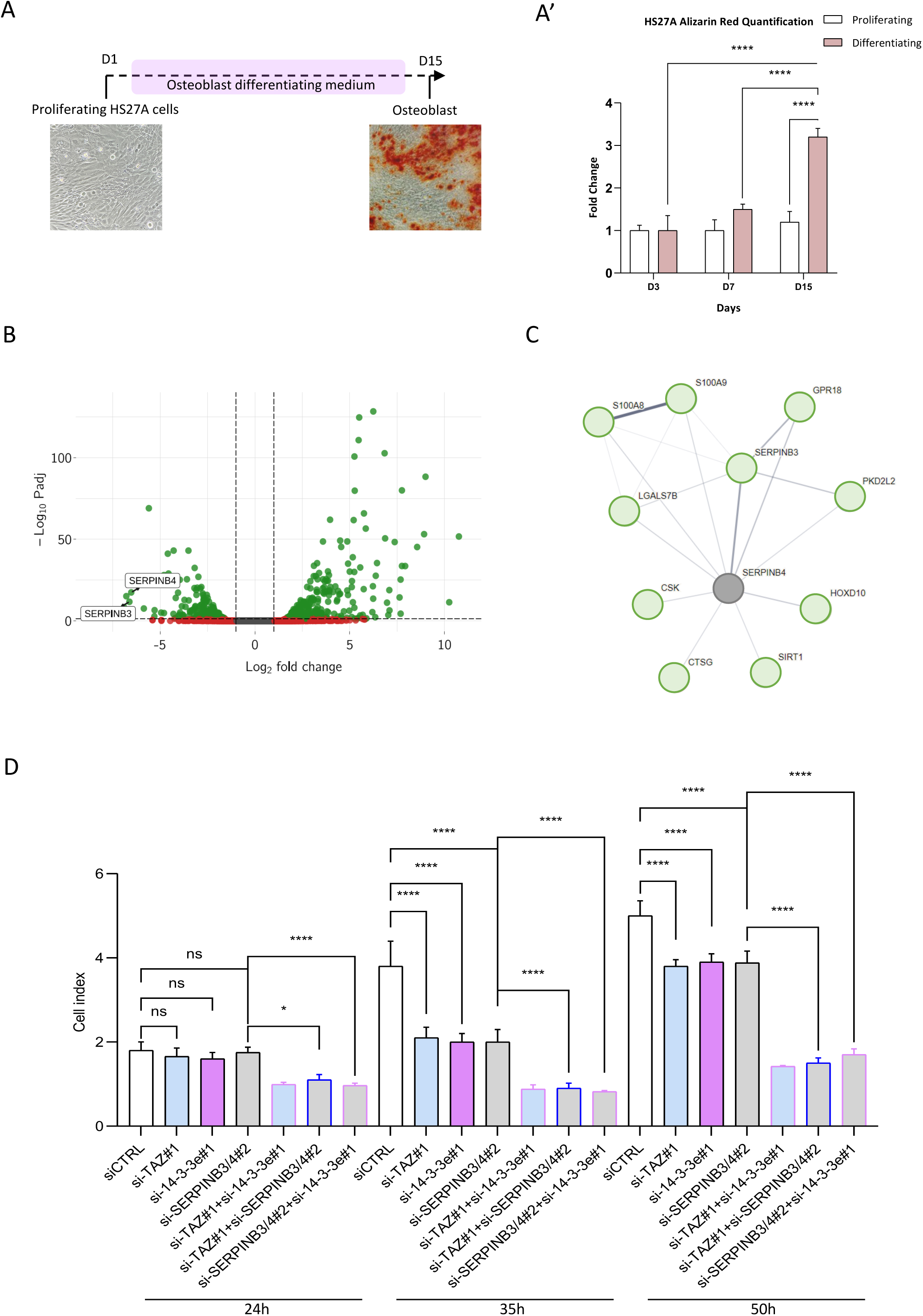
Role of SERPINB3 and SERPINB4 in TAZ/14-3-3e-dependent proliferation of HS27A mesenchymal stem cells. **(A)** HS27A cells differentiate into osteoblast and are stained with alizarin (red) after 15 days of culture in an osteoblast differentiation medium. **(A’)** Statistical quantification of the HS27A cell alizarin staining at day 3, 7 and 15 of culture. Statistical tests were performed with one-way ANOVA (\*\*\*\**p* < 0.0001). **(B)** Distribution as a volcano plot of up-(Log2FC>0.5 and Log10*pvalue*>0,8) or down-(Log2FC<-0.5 and Log10*pvalue*>-0,8) regulated genes during the differentiation of HS27A cells (as deduced from RNA-seq. experiments performed at day 1 and day 15). **(C)** String interactome for SERPINB4 (based on 0.15 interaction score of experimental evidence and database and pathway co-occurrence). **(D)** Quantification with xCELLigence assays of HS27A cells transfected with the different siRNAs, as indicated. siC = siRNA control (see also Materials and Methods). Measures were performed at different time points post-transfection and resulted from three independent biological replicates. Two-way ANOVA with Tukey’s multiple comparisons; * *p* < 0.05 and **** *p* < 0.0001.

To test this hypothesis, we generated two different siRNAs targeting *TAZ*, *14-3-3e* or *SERPINB3* and *B4*. The specificity of each siRNA was individually tested in HS27A cells (Fig. S9 and Materials and Methods) and we next assessed their effect on HS27A cells proliferation, either alone or in various heterologous combinations. Results showed that the individual targeting of *TAZ, SERPINB3/B4* or *14-3-3e* affected the proliferation rate of HS27A cells at 35h post transfection, and this effect tended to fade over time (at 50h, which could be explained by siRNA dilution over cell generations; Fig. 4D). Importantly, combining siRNAs against *SERPINB3/B4* and *TAZ* or *SERPINB3/B4* and *14-3-3e* already impacted proliferation at 24h post-transfection, and further decreased HS27A proliferation rate when compared to individual siRNAs at 50h (Fig. 4D). These combinatorial effects were similar to the one observed when combining *TAZ* and *14-3-3* siRNAs and were more persistent over time when compared to individual siRNAs (Fig. 4D). These results show that SERPINB3 and B4 are required for the proliferation of HS27A cells and act in concert with TAZ and 14-3-3e. This effect was independent of cross-regulatory relationships given that the knock-down of *SERPINB3* and *B4* had no effect on *TAZ* or *14-3-3e* expression (Fig. S9). We propose that the combinatorial effect of siRNAs could result in a more complete loss of functional protein complex assemblies given that each siRNA was not 100% effective (Fig. S9). In this context, *SERPINB3* and *B4* could have a redundant role for promoting the proliferative activity of TAZ and 14-3-3e through the formation of specific trimeric protein complexes.

### Specific interaction interfaces allow the formation of trimeric TAZ/14-3-3e/SERPBIN4 complexes

To understand how SERPINB3 or B4 could form a trimeric complex with TAZ and 14-3-3e, we used the artificial intelligence (AI) platform AlphaFold-multimer to predict hetero-oligomeric structures (https://alphafold.ebi.ac.uk/ and [37]), and the webserver ENDscript to analyze intermolecular contacts [38]. Although the top-ranked predicted model for SERPINB3 and B4 had a relatively low interface predicted template modelling (ipTM) score (0,4), as it can be expected because of long disordered regions of TAZ, they both showed the emplacement of a disordered 306-322 C-terminal segment of TAZ in between 14-3-3e and SERPINB3 or B4 (Fig. S10-S11). This segment of TAZ contains a short linear interaction motif (SLIM) that could be phosphorylated by LATS kinases ([39–41] and Fig. S10). Importantly, this SLIM is the only contacting region of TAZ with SERPINB3 or B4 (Fig. S10-11). In particular, a unique residue, Glu309, is depicted to contact two Lys residues of SERPINB4 in the trimeric complex with 14-3-3e (Fig. S10 and S11). In contrast, several residues of the SLIM are establishing contacts with 14-3-3e in the same trimeric protein complex (Fig. S10-11). Overall, these predictions highlight that the C-terminal SLIM of TAZ could constitute a central node for making strong interactions with both SERPINB3 or B4 and 14-3-3e in the trimeric complex.

To further confirm these predictions, we took advantage of bicolor BiFC to analyze interaction properties between TAZ, 14-3-3e and SERPINB4 in a live cell context. As described previously (Fig. 2B), bicolor BiFC was achieved by co-expressing VN-TAZ and CC-14-3-3e together CN-SERPINB4 to look at the Venus-like (VN-TAZ/CC-14-3-3e) and Cerulean-like (CN-SERPB3-4/CC-14-3-3e) fluorescent signals in living HEK293T cells (Fig. 5A). Results showed identical diffuse profiles with few enrichment dots in the cytoplasm for the two BiFC signals, demonstrating that TAZ-14-3-3e and 14-3-3e-SERPINB4 interactions occurred at the same places and could therefore be part of a same protein complex (Fig. 5A’). To further confirm that SERPINB4 could interact with both TAZ and 14-3-3e, we performed bicolor BiFC by expressing CC-SERPINB4 together with VN-TAZ and CN-14-3-3e, allowing looking at TAZ-SERPINB4 and 14-3-3e-SERPINB4 interactions in green and cyan, respectively (Fig. 5B). Again, the two BiFC profiles were superposable in the cytoplasm (Fig. 5B’), indicating that the two interactions occurred at the same locations within the cell. Altogether, bicolor BiFC observations are in agreement with the potential formation of trimeric TAZ/14-3-3e/SERPINB4 complexes.

**Figure 5:**
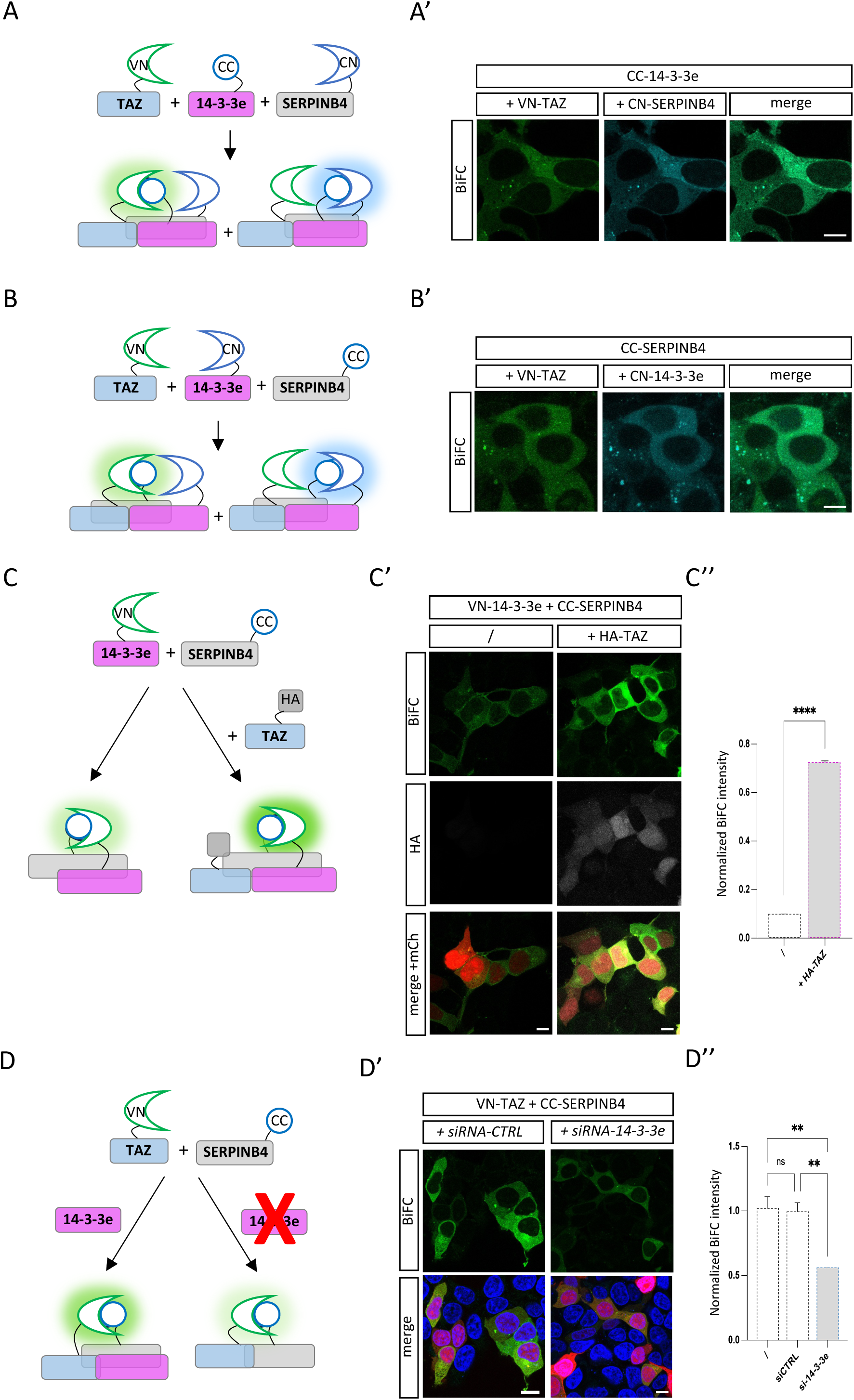
TAZ, 14-3-3e and SERPINB4 establish cooperative protein-protein interactions in living HEK293T cells. **(A)** Scheme of bicolor BiFC illustrating Venus-and Cerulean-like signals resulting from TAZ/14-3-3e and 14-3-3e/SERPINB4 interactions, respectively. **(A’)** Illustrative confocal acquisition showing the simultaneous interaction profile of TAZ/14-3-3e (green) and 14-3-3e/SERPINB4 (blue) in living HEK293T cells. **(B)** Scheme of bicolor BiFC illustrating Venus-and Cerulean-like signals resulting from TAZ/SERPINB4 and 14-3-3e/SERPINB4 interactions, respectively. **(B’)** Illustrative confocal acquisition showing the simultaneous interaction profile of TAZ/SERPINB4 (green) and 14-3-3e/SERPINB4 (blue) in living HEK293T cells. **(C)** Scheme of BiFC performed between 14-3-3e and SERPINB4 (green) when co-expressing or not HA-tagged TAZ. **(C’)** Illustrative confocal acquisition of BiFC (green) between 14-3-3e and SERPINB4 without (first column) or with HA-TAZ overexpression, as indicated. HA-TAZ was revealed with an anti-HA antibody (gray) and mCherry (red) stains for transfection efficiency. **(C’’)** Quantification of the BiFC (normalized over transfection efficiency) between 14-3-3e and SERPINB4 in the absence (first column) or presence (second column) of HA-TAZ. **(D)** Scheme of BiFC performed between TAZ and SERPINB4 (green) when affecting or not the expression of endogenous *14-3-3e*. **(D’)** Illustrative confocal acquisition of BiFC between TAZ and SERPINB4 (green) in the siRNA control (ctrl) condition or when co-transfecting siRNA targeting endogenous *14-3-3e*, as indicated. Merge with Hoechst (blue) and mCherry (red) stains for nuclei and transfection efficiency, respectively. **(D’’)** Quantification of the BiFC (normalized over transfection efficiency) between TAZ and SERPINB4 with no siRNA (first column), control siRNA (second column) or siRNA against *14-3-3e* (third column). Statistical tests were performed with one-way ANOVA from three biological replicates (ns: non-significant, \*\*\**p* < 0.001, \*\**p* < 0.01). Scale bar = 20 µm.

To assess whether the formation of these trimeric complexes could rely on cooperative interactions between the three partners, we took advantage of BiFC as a simple readout of interaction affinity. More precisely, we first expressed only the VN-14-3-3e and CC-SERPINB4 constructs, which led to Venus-like fluorescent signal (Fig. 5C-C’). We reasoned that these BiFC complexes could potentially be established for their large majority without endogenous TAZ given that BiFC constructs were overexpressed with the dox-inducible promoter and that *TAZ* was weakly expressed in HEK293T cells (as deduced from RNA-seq experiments, Kundlacz et al., in preparation). Accordingly, co-overexpressing a HA-tagged version of TAZ together with VN-14-3-3e and CC-SERPINB4 led to a strong increase of BiFC signals (around 7 folds, Fig. 5C-C”). This observation demonstrates that the formation of a trimeric complex with TAZ promotes the interaction between 14-3-3e and SERPINB4.

To also analyze the contribution of 14-3-3e on TAZ-SERPINB4 interaction, we performed BiFC between VN-TAZ and CC-SERPINB4 when affecting endogenous *14-3-3e* (Fig. 5D), which is more strongly expressed than *TAZ* in HEK293T cells (Kundlacz et al., in preparation). We observed that the depletion of endogenous *14-3-3e* (around 60% loss was obtained upon siRNA transfection, S9) led to strong decrease of BiFC signals (around 50% loss, Fig. 5D’-D”). Thus, we concluded that endogenous 14-3-3e was sufficiently highly expressed to participate to the BiFC complexes and promote TAZ-SERPINB4 interaction.

Altogether, these experiments established that binary interactions between SERPINB4 and TAZ or SERPINB4 and 14-3-3e were increased in the presence of the third partner, strongly suggesting that TAZ, 14-3-3e and SERPINB4 could establish cooperative interactions to trimeric protein complex.

To experimentally validate the predictive interaction interface of TAZ with SERPINB4, we mutated the residue Glu309 into an Alanine residue (construct TAZ^E309A^), anticipating that this mutation could affect TAZ-SERPINB4 interaction specifically in the context of TAZ/SERPINB4/14-3-3e trimeric complex. Indeed, interaction interfaces are all predicted to be different when comparing the trimeric complex to the different dimeric complexes (Fig. S12-14). Of interest, the residue Glu309 of TAZ is not predicted to contact any SERPINB4 residues in the dimeric TAZ/SERPINB4 complex (Fig. S12). These predictions underscore that binary interactions could be completely remodeled in the presence of the third partner.

The correct expression and localization of TAZ^E309A^ was verified (Fig. S10) and the effect of the TAZ punctual mutation was analyzed by doing bicolor BiFC to simultaneously analyze the interaction of SERPINB4 with either TAZ^E309A^ or 14-3-3e (Fig. 6A). Results showed 80% loss of BiFC between SERPINB4 and TAZ^E309A^, thus confirming that the residue Glu309 of TAZ was central for recruiting SERPINB4 in the context of the trimeric complex (Fig. 6A’-A”). The inability of TAZ^E309A^ to interact with SERPINB4 was also indirectly demonstrated by the absence of any promoting effect of HA-TAZ^E309A^ when over-expressed with VN-14-3-3e and CC-SERPINB4 (Fig. 6B-B”), indicating that HA-TAZ^E309A^ lost most of its ability to form a trimeric complex with SERPINB4 and 14-3-3e.

**Figure 6:**
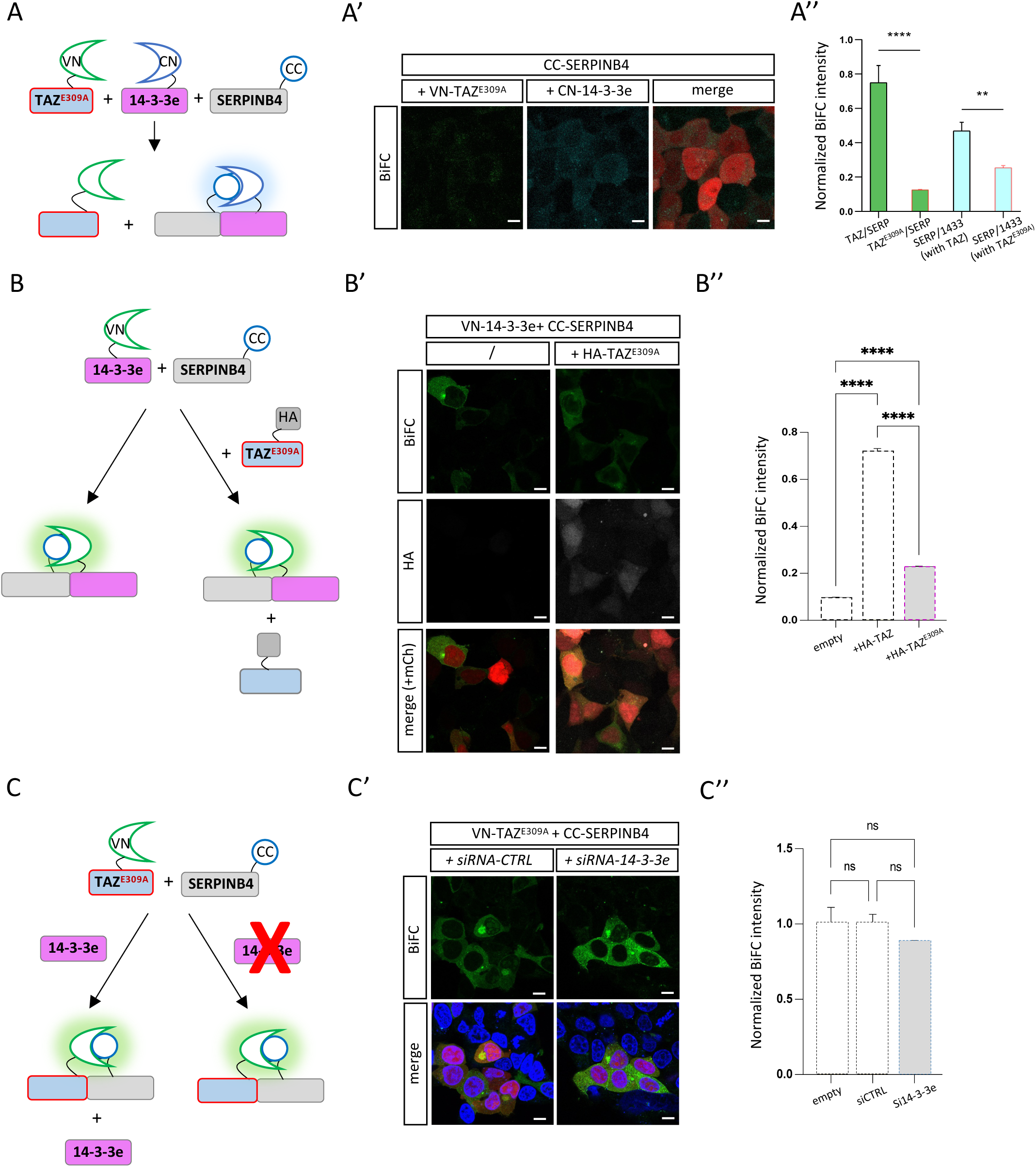
The Glu309 residue of TAZ is critical for recruiting SERPINB4 specifically in the presence of 14-3-3e. **(A)** Scheme of bicolor BiFC with the mutated TAZ^E309A^ construct and illustrating the confocal observations in A’. **(A’)** Illustrative confocal acquisition showing the loss of BiFC between mutated TAZ^E309A^ and SERPINB4 (green) and the decreased BiFC between SERPINB4 and 14-3-3e (blue) in living HEK293T cells. mCherry (red) stains for transfection efficiency in the merge picture. **(A’’)** Quantification of the BiFC (normalized over transfection efficiency) between wild type or mutated TAZ and SERPINB4 (first two columns), and between SERPINB4 and 14-3-3e with wild type or mutated TAZ (last two columns) co-transfection conditions. **(B)** Scheme of BiFC performed between 14-3-3e and SERPINB4 (green) when co-expressing or not the HA-tagged mutated form of TAZ. **(B’)** Illustrative confocal acquisition of BiFC between 14-3-3e and SERPINB4 without (first column) or with (second column) co-expression of mutated HA-TAZ^E309A^, as indicated. Mutated HA-TAZ^E309A^ was revealed with an anti-HA antibody (gray). mCherry (red) stains for transfection efficiency. **(B’’)** Quantification of BiFC (normalized over transfection efficiency) between 14-3-3e and SERPINB4 in the absence (first column) or presence of wild type HA-TAZ (second column) or mutated HA-TAZ^E309A^ (third column). **(C)** Scheme of BiFC performed between mutated TAZ^E309A^ and SERPINB4 (green) when affecting or not the expression of endogenous *14-3-3e*. **(C’)** Illustrative confocal acquisition of BiFC between mutated TAZ^E309A^ and SERPINB4 (green) in the siRNA control (ctrl) condition or when co-transfecting siRNA targeting endogenous *14-3-3e*, as indicated. Merge with Hoechst (blue) and mCherry (green) stains for nuclei and transfection efficiency, respectively. **(C’’)** Quantification of the BiFC (normalized over transfection efficiency) between mutated TAZ^E309A^ and SERPINB4 with no siRNA (first column), control siRNA (second column) or siRNA against *14-3-3e* (third column). Statistical tests were performed with one-way ANOVA from three biological replicates (ns: non-significant, \*\**p* < 0.01, \*\*\*\**p* < 0.0001). Scale bar = 20 µm.

Surprisingly, bicolor BiFC in the context of VN-TAZ^E309A^ also showed a strong decrease (around 50%) of signal between CN-14-3-3e and CC-SERPINB4 (Fig. 6A’-A”). Thus, the loss of interaction between TAZ^E309A^ and SERPINB4 also affected the interaction between SERPINB4 and 14-3-3e, underlining that the interaction interfaces between these two proteins was remodeled (and with a lower affinity) when TAZ was not interacting with SERPINB4.

To further confirm that interaction interfaces are different when considering the trimeric and dimeric complexes, we analyzed the effect of the mutation of the Glu309 residue of TAZ in the absence of 14-3-3e. To this end, we repeated the analysis of BiFC between VN-TAZ^E309A^ with CC-SERPINB4, in wild type condition or by simultaneously expressing *14-3-3e* siRNA to affect endogenous 14-3-3e (Fig. 6D). First, we observed BiFC despite the presence of endogenous *14-3-3e* in wild type condition (Fig. 6D’). This observation confirmed that VN-TAZ^E309A^/CC-SERPINB4 dimeric complexes could be formed in excess with regard to endogenous 14-3-3e. Importantly, depleting endogenous *14-3-3e* did not affect the BiFC signal intensity, demonstrating that TAZ^E309A^/SERPINB4 complexes were formed through distinct and 14-3-3e-independent interaction interfaces as compared to TAZ-SERPINB4 interactions established in the context of trimeric TAZ/SERPINB4/14-3-3e BiFC complexes (Fig. 6D’-D”).

Collectively, these experiments confirmed that the Glu309 residue of TAZ is critical for recruiting SERPINB4 in the specific context of the trimeric association with 14-3-3e. In addition, this Glu309-dependent interaction allows SERPINB4 to strongly interact with 14-3-3e, thus showing the mutual dependency of each partner to form a cooperative trimeric protein complex.

## Discussion

In this study, we developed and validated Bi-nano-ID, a novel method that combines BiFC with proximity-dependent biotinylation, to investigate binary protein interactions and their associated interactomes in living cells. By using TAZ/14-3-3e and TAZ/TEAD2 interactions as a proof-of-concept, we demonstrated the efficiency of Bi-nano-ID in capturing and characterizing these interactions within their specific cellular compartments.

### Advantages of Bi-nano-ID over other experimental approaches and future potential improvements

To date, the unique alternative methodologies to capture interactomes of dimeric complexes rely on split-BioID and split-TurboID enzymes, which can be reconstituted upon spatial proximity. These approaches have been used to identify interactomes of different dimeric protein complexes and to monitor [8,42,43] or capture interactions in specific cellular compartments [5]. However, these systems require long biotin incubation times (minimum 8 hours) due to the decreased activity of the reconstituted TurboID. In contrast, Bi-nano-ID integrates BiFC, which is well established for providing a specific readout of many different types of PPIs, allowing visualizing and validating the correct interaction profile of the dimeric protein complex in living cells. In addition, the use of a BiFC-recognyzing nanobody fused to full length TurboID enables to exploit the robust and rapid biotinylation capabilities of full length TurboID with shorter time of biotin incubation when compared to split-TurboID systems. Incubation times applied with Bi-nano-ID are still in the range of few hours, but this time could potentially be diminished if it is a critical parameter. For example, in our work we decided for 180min incubation time, but a significant enrichment (also to a less extent) could already be obtained at 90min incubation time (Fig. S4). Getting access to the fine temporal dynamics (range of minutes) of a binary complex interactome will require developing tools with other complementation systems that are reversible (unlike BiFC, see for example [44]). In any case, the Nanobody-TurboID combination constitutes the most sensitive tool for detecting interactions of binary protein complexes within defined subcellular localizations. Finally, by using fragments compatible for bicolor BiFC, Bi-nano-ID allows to rapidly validate candidate proteins of the interactome together with the bait dimeric complex in living cells. In addition, human gene libraires have also been generated in fusion with the CC fragment to perform large-scale BiFC interaction screens in human living cells (*Kundlacz et al. in preparation* and [45]). In this context, Bi-nano-ID could help to further characterize the endogenous interactome of the VN-bait protein associated to a candidate CC-cofactor identified from a BiFC interaction screen.

### Bi-nano-ID identified distinct interaction partners for TAZ/14-3-3e and TAZ/TEAD2 complexes

Interactions between TAZ and 14-3-3e or TAZ and TEAD2 were visualized using BiFC, showing distinct cytoplasmic and nuclear fluorescence patterns respectively, and confirming their subcellular localization as influenced by the Hippo signaling pathway. This initial validation set the foundation for integrating the csGBP nanobody fused with the TurboID enzyme, which led to the biotinylation of proteins in proximity to the BiFC complexes. The subsequent biotinylation was specific and colocalized with the BiFC signals, as revealed by immunofluorescence and western blot analyses, indicating the precision and reliability of Bi-nano-ID in live-cell contexts.

Due to the weak recognition of the VN173 fragment by the csGBP nanobody, we also tested our tools when using VN155-TAZ, which was not recognized by the nanobody in previous work [19]. These assays showed that BiFC was weak and not exploitable with 14-3-3e or TEAD2 fused to the CC fragment. Consequently, biotinylation was also less efficient. We concluded that VN155 was much less appropriate than VN173 in our bicolor complementation strategy, which was also consistent with previous work [45]. In addition, using VN173-TAZ alone as a negative control was efficient for discriminating the background resulting from the weak affinity of the csGBP nanobody for the VN173 fragment in the mass spectrometry analysis.

Our work identified a substantial number of proteins in proximity to TAZ/14-3-3e and TAZ/TEAD2 complexes. The stringent filtering process ensured high specificity, leading to the identification of 96 candidates for TAZ/14-3-3e and 42 candidates for TAZ/TEAD2. Further refinement excluded proteins identified with TAZ-TurboID, which were considered as generic interaction partners when compared to interactions that could be established with BiFC-stabilized TAZ/14-3-3e and TAZ/TEAD2 complexes. This filtering step led to the identification of 49 TAZ/14-3-3e interactors and 29 TAZ/TEAD2 interactors. 17 interactions were common to the two complexes, demonstrating that Bi-nano-ID was successful in capturing interactions that were specific of TAZ/14-3-3e (32 interactions) and TAZ/TEAD2 (12 interactions). Gene ontology enrichment analyses revealed that TAZ-TurboID, TAZ/14-3-3e and TAZ-TEAD2 interactors all predominantly participate in Hippo pathway regulation, highlighting that these novel interactions could modulate TAZ function. Importantly, TAZ/14-3-3e and TAZ/TEAD2 interactomes displayed clear distinct GO term enrichments, allowing assigning specific functions of TAZ depending on its association with either 14-3-3e or TEAD2 cofactors. Collectively, these results demonstrate the suitability of Bi-nano-ID for capturing and distinguishing interactomes of a bait protein associated with a different cofactor in two different cellular compartments.

### Molecular insights into TAZ/14-3-3e/SERPINB4 complex formation and its biological implications

Our detailed molecular dissection using bicolor BiFC and mutagenesis identified a critical interaction interface for both 14-3-3e and SERPINB4 within the TAZ protein. More particularly, this region encompasses residues 306-322 and is predicted to be phosphorylated by LATs kinases, therefore under dynamic regulation depending on the activation of the Hippo pathway. Importantly, mutating the single Glu309 residue of TAZ predicted to form electrostatic interactions with two Lys residues of SERPINB4 was sufficient to abolish all cooperative interactions involved in the trimeric complex assembly. In particular, the Glu309Ala mutation was able to also affect the interaction between 14-3-3e and SERPINB4 but only in the context of the trimeric complex, highlighting that 14-3-3e and SERPINB4 have TAZ-dependent and TAZ-independent interaction modes (Fig. 7). Similar interaction properties could apply for TAZ/14-3-3e/SERPINB3 complexes although it remains to be firmly demonstrated. We suggest that the TAZ-dependent mode of interaction, therefore interaction in the context of trimeric TAZ/14-3-3e/SERPINB3/4 complexes, could be more stable and potentiate the proliferative activity of TAZ/14-3-3 complexes (Fig. 7). This hypothesis results from our assays in human bone marrow stromal cells (HS27A) but could potentially apply more generally in cell progression, proliferation and metastasis for cancer biology and tissue regeneration. In this context, the identification of the most relevant interaction interface used in the trimeric and not the dimeric complex could have strong incidence for future therapeutic development.

**Figure 7.**
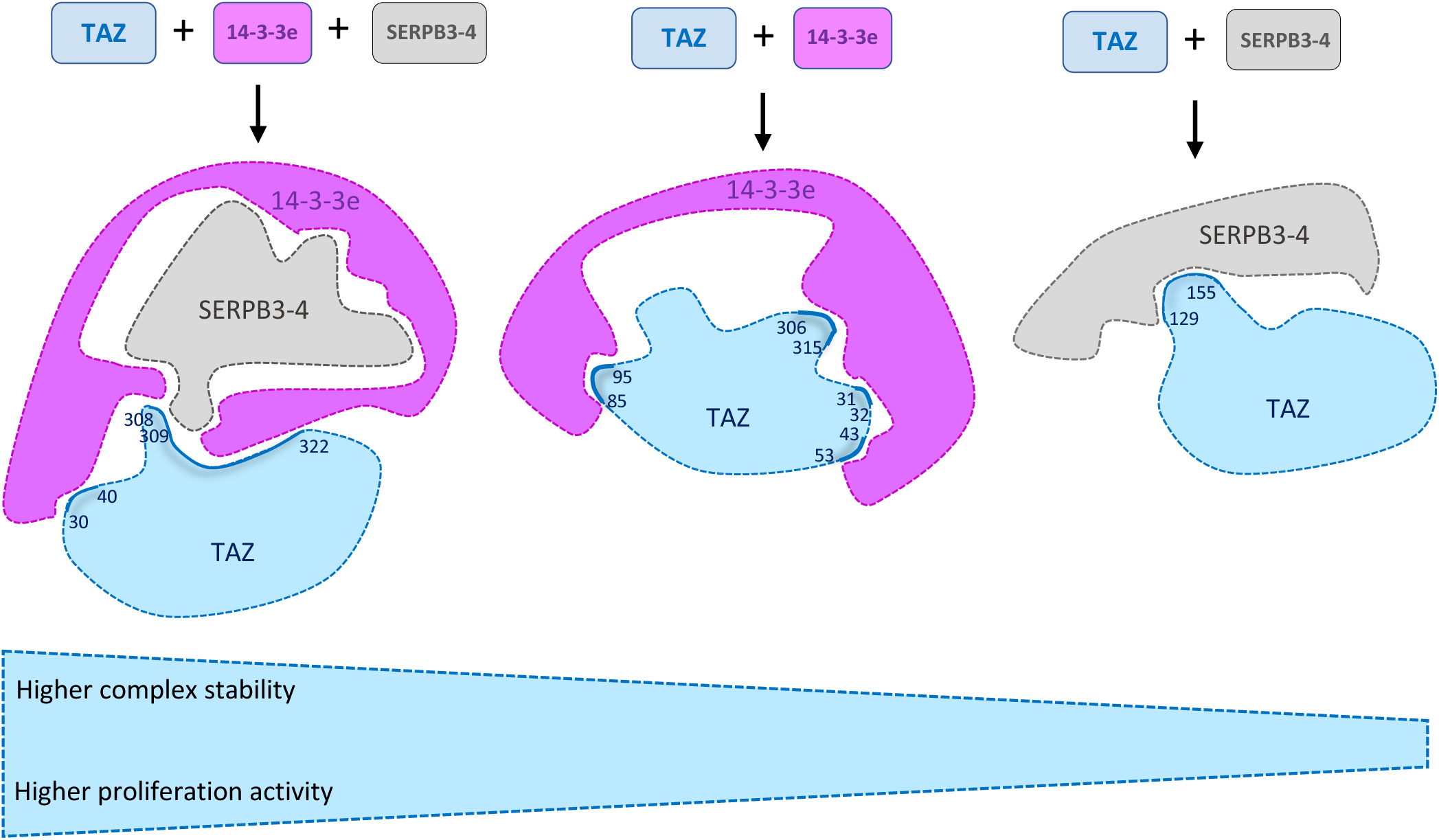
A specific interaction mode underlies the formation of TAZ/14-3-3e/SERPINB4 trimeric complexes. The Glu309 of TAZ is a unique contact residue for SERPINB4, being responsible of the formation of a cooperative trimeric complex with 14-3-3e. The trimeric TAZ/14-3-3e/SERPINB4 complex relies on specific interaction interfaces that are not used in the dimeric complexes. Accordingly, the TAZ*^E309A^* mutated form can interact with SERPINB4 and the corresponding dimeric complex is insensitive to the absence of 14-3-3e. Thus, the formation of trimeric TAZ/14-3-3e/SERPINB4 complexes relies on strong protein interaction remodeling when compared to the dimeric TAZ/14-3-3e or TAZ/SERPINB4 complexes. This remodeling could have incidence on the overall protein complex stability and proliferation activity. Sites of contact with 14-3-3e and SERPINB4 are highlighted in TAZ for the dimeric and trimeric complexes (as deduced from AlphaFold and Endscript predictions).

In conclusion, our work highlights the importance of considering interactomes of dimeric protein complexes over single bait proteins, and the relevance of Bi-nano-ID in this perspective.

## Materials and Methods

### Cell culture

HEK293T cells were purchased from the European Collection of Authenticated Cell Cultures (ECACC) through the biological resource center CelluloNet (AniRA platform of the UAR3444, Lyon, France). Human bone marrow stromal cells (HS27A) were a gift from the team of Dr. Véronique Maguer-Satta (CRCL). HEK-293T cells were cultured as a monolayer in Dulbecco’s modified Eagles medium (DMEM-GlutaMAX-I, Gibco by Life Technologies) supplemented with 10% (*v/v*) heat-inactivated fetal bovine serum (FBS) (Gibco by Life Technologies), 1% (*v/v*) Penicillin-Streptomycin (5,000U penicillin and 5mg streptomycin/mL, Gibco), and 5 ml of 100 mM sodium pyruvate (Gibco by Life Technologies) incubating at 37°C, in an atmosphere of 5 % CO_2_. HS27A cells were cultured in RPMI 1640 (Gibco by Life Technologies) supplemented with 10% (*v*/*v*) heat-inactivated fetal bovine serum (FBS) (Gibco by Life Technologies) and 1% (*v/v*) Penicillin-Streptomycin (5,000U penicillin and 5mg streptomycin/mL, Gibco) incubating at 37°C, in an atmosphere of 5 % CO2.

### Cloning

Plasmids used in this study were cloned either by gateway cloning or conventional restriction digestion into pLIX _403 vector (a gift from David Root; Addgene plasmid # 41395; http://n2t.net/addgene:41395, RRID: Addgene_41395). All constructs are placed under the same tet-off dox-inducible promoter. All constructs were sequenced (Genewiz company, Germany) before using. Oligonucleotides used for the cloning are listed (Table S4).

### Fluorescent immunostaining

50.10^5^ HEK-293T cells were seeded on glass coverslips in 24-well plate for 18-24 hours to about 80% confluency, then transiently transfected with 250 ng of each plasmid using jetPRIME (Polyplus, Ref 114-15) and following manufacturer’s instructions in the presence of doxycycline (dox, 100 ng/ml). After 14h of dox induction, cells were washed with PBS and replaced by fresh complete media without dox for 6h at 37°C. Cells were then cultured in 50 µM Biotin for 3h at 37°C under 5% CO_2_. Cells were then washed twice with 1X PBS and fixed with 4% (*v/v*) paraformaldehyde at 4°C for 15 min. In between each step, cells were washed with 1X PBS. Cells were permeabilized with 0,1% Triton for 10 min followed by a blocking step in 2% (*w/v*) BSA for 1h at RT. Cells were then incubated with primary antibody (Rabbit-anti-GFP A11122 Invitrogen at 1/1000) in PBS supplemented with 2% (w/v) BSA for 3 hr at RT. Next, samples were incubated with anti-rabbit-Alexa Fluor 488 and Alexa Fluor 555-Streptavidin to reveal biotinylated proteins, in PBS supplemented with 2% (w/v) BSA for 1 hr at RT. Finally, cell-coated coverslips were mounted in VECTASHIELD antifade mounting medium with DAPI (VECTOR, Cat No. LS-J1033-10) and imaged by confocal microscopy (Zeiss LSM780).

### Western blots

Cells were cultured in Biotin (50 µM) at 37°C under 5% CO_2_ at several incubation times (10 min, 90min, 180min, and 240min). Cells were washed twice with 10 ml ice-cold DPBS, harvested by scraping, pelleted by centrifugation at 1,400 r.p.m. for 3 min, and either processed immediately or stored at -80°C before further analysis.

The protocol of TurboID labeling (with TurboID-HA or TAZ-TurboID-HA constructs) is similar to the approach followed for Bi-nano-ID labeling, except that cells were incubated with biotin 50 µM for 10 min.

Collected cell pellets for both Western blotting (WB) and proteomic analysis were resuspended in ice-cold RIPA/1%SDS-cell lysis buffer (50mM Tris pH8, 150mM NaCl, 0.5% sodium deoxycholate, 1% NP40/IGEPAL) (volume ratio between cell pellet: buffer = 1:3) and incubated on ice for 20-30 min, maintaining constant vortex (2-3 times). The cells were then treated with Benzonase Nuclease (Sigma, E1014-25KU) for 10 to 20 min on ice or to observe the solution becomes less viscous. This was subjected to spin at 14000×g at 4°C for 10 min. The supernatant was then transferred to 1,5 Eppendorf tube. To test the expression of the constructs, 10% of the protein extracts were kept for WB as input fraction. Proteins were quantified using Pierce BCA Protein Assay (Thermofischer, A55865). Streptavidin magnetic beads (Sigma, 88817) were equilibrated with two PBS washes and resuspended in RIPA buffer supplemented with 1% SDS (50 mM Tris pH 8, 150 mM NaCl, 0.5% sodium deoxycholate, 1% NP40, 1% SDS). 1 mg of protein extracts were incubated with 40 µl of streptavidin beads in a final volume of 1 ml RIPA-SDS overnight at 4°C on a rotating wheel. Beads were then washed twice with SDS-Buffer (10 mM Tris.HCl, 1 mM EDTA, 1% SDS, 200 mM NaCl), twice with RIPA-SDS and twice with acetonitrile buffer (20% acetonitrile in MS-grade water). For WB validation, acetonitrile washing step was skipped.

For western blot analysis, proteins (Input and AP) were resolved on 12% SDS-PAGE, blotted onto PVDF membrane (Biorad) and probed with specific antibodies after saturation. The antibodies (and their dilution) used in this study were, Histone 3 (1791 Abcam, 1/10,000), GFP (A11122 Life Technologies, 1/1000), and Streptavidin-HRP (SA10001 Invitogen, 1/5000). Developing was performed using chemiluminescence reaction (ECL, Cytiva) with secondary antibody coupled to HRP (Promega, 1/5000).

### Mass spectrometry preparation

Streptavidin beads were resuspended in 14 µl of ammonium bicarbonate 50 mM and proteins were subjected to reduction with 1 µl DTT (100 µM, 57°C, 45min) and alkylation with 1µl iodoacetamide (500µM, RT, 45min at dark). The proteins were then digested with a mixture of endoproteinase Lys-C/trypsin (ratio protein/enzyme 1/100, overnight, 37°C, 500 rpm). After a cleaning step with SP3 beads, peptides were dried, suspended in Formic Acid (FA) 0.1%, DMSO 2% and injected in the mass spectrometer. Samples were analyzed on a Q Exactive HF mass spectrometer coupled with a nanoRSLC Ultimate 3000 system (ThermoFisher Scientific). Peptides samples were loaded on a C18 Acclaim PepMap100 trap-column 300 µm ID x 5 mm, 5 µm, 100Å (ThermoFisher Scientific) and separated on a C18 Acclaim Pepmap100 nano-column, 50 cm x 75 µm i.d, 2 µm, 100 Å (ThermoFisher Scientific) with a 60 minutes linear gradient from 3% to 90% buffer B (A: 0.1% FA in H_2_O, B: 0.1% FA in ACN). The flow rate was maintained at 300 nL/min and the oven temperature was kept constant at 40°C. Peptides were analysed using a DDA TOP20 HCD method: MS data were acquired in a data dependent strategy (DDA) selecting the fragmentation events based on the 20 most abundant precursor ions in a survey scan (350-1400 Th). Resolutions of the survey and MS/MS scans were respectively set at 60,000 and 15,000 at m/z 200 Th. The Ion Target Values for the survey and the MS/MS scans in the Orbitrap were set to 3E6 and 1E5 respectively and the maximum injection times were set to 60 ms for both MS and MS/MS scans. Parameters for acquiring HCD MS/MS spectra were as follows: collision energy = 27 and isolation window = 2 m/z. The precursors with unknown charge state, charge state of 1 and 8 or greater than 8 were excluded. Peptides selected for MS/MS acquisition were then placed on an exclusion list for 30 s using the dynamic exclusion mode to limit duplicate spectra.

### Mass spectrometry analyses

Each batch of experiment included TurboID and TAZ fused to TurboID samples (Table S2), and csGBP-TurboID, csGBP-TurboID with VN-TAZ, csGBP-TurboID with VN-TAZ and CC-14-3-3e, csGBP-TurboID with VN-TAZ and CC-TEAD2 samples (Table S3). Each condition was performed in three independent biological replicates. Raw mass spectrometry data were analysed using MaxQuant free software including the Andromeda search Engine [46]. Using *Homo sapiens* (Swissprot) database peptide were identified. Default parameters of MaxQuant were used with the following modifications: digestion by Trypsin/P and LysC, lysine biotinylation as variable modification (as well as methionine oxidation and N-terminal acetylation), cytosine carbamidomethylation as fixed modification, Instrument set Orbitrap (with precursor tolerance 20 ppm, MS tolerance 0.5 Da), match between runs option was activated, FDR 1%, label-free quantification (LFQ) and iBAQ were calculated (Tables S2 and S3). Protein enrichment was calculated using the LFQ Log2 ratio and normalized on the median value (Table S2 and S3). For each ratio, distribution (90%) and corresponding standard deviations (SD) were calculated to define the proteins significantly enriched (ratio > confidence interval defined as median ± 2 SD). Imputation of value divided by 0 (referred to infinite) has been performed for confidence intervals calculation (Table S1 and S2). Subsequently, proteins significantly enriched in at least 2 replicates were considered biologically relevant considering biological variability and stochasticity of the MS-process and used for further analysis (Table S2 and S3). Enriched proteins from the different samples were then compared with discriminate proteins enriched in proximity with csGBP-TurboID individually or in proximity with VN173-TAZ alone. Details about the filtering process are detailed (Fig. S5). For TAZ generic candidates, protein enrichment was calculated using the LFQ Log2 ratio of TAZ-TurboID/TurboID and normalized on the median value with distribution (90%) and corresponding standard deviations (SD) were calculated to define the proteins significantly enriched (ratio > confidence interval defined as median ± 2 SD).

The mass spectrometry proteomics data have been deposited to the Center for Computational Mass Spectrometry repository (University of California, San Diego) via the MassIVE tool with the dataset identified MassIVE MSV000095990.

### Data analysis and visualisation

For proteome analysis, Perseus free software was used to generate clustering visualization (PCA, [47]), based on LFQ log10 value of protein expressed after Perseus canonical filtering (Reverse, Potential Contaminant, only identified by site) and replacement of missing values. Functional networks of TAZ interactome were generated with STRING software [48], based on 0.15 interaction score of experimental evidence and database and pathway co-occurrence. Visualization of networks was built with Cytoscape free software [49]. For GO-Term annotations and over-represented GO-Term related to biological process analysis was performed using PANTHER [50] and represented using Prism. The selection of significant GO-term was corrected by FDR<0.05, a fold enrichment higher than 10 and a raw p-value>0.05. Statistical analyses were performed using one-way ANOVA (BiFC quantification, siRNA effect, proliferation assay). Predictions of TAZ/1433e/SERPINB4 interaction model was performed using Alphafold [37].

### BiFC and bicolor BiFC validation

For transfection, 3.10^5^ HEK293T cells were seeded on glass coverslips in 6-well plates and incubated for 24h. Then, cells were transfected with jetPRIME (Ref 114-15, Polyplus Transfection, France) following manufacturer’s instructions. A total of 1 µg of plasmid DNA (CC, VN and CC tagged proteins constructs including mcherry reporter to assess transfection efficiency) was transfected per well. After 14h of incubation in the presence of doxycycline (100 ng/mL), the cell-coated coverslip was taken and mounted carefully on a glass slide for image capturing under Zeiss LSM780 confocal microscope (Carl Zeiss, Jena, Germany). All samples were imaged using identical settings and were quantified as previously described [45]. Three biological replicates were systematically performed.

### RTqPCR and siRNA

For siRNA transfection, 3.10^5^ cells per ml were seeded on 6-well plates, and 10 pmol/mL of total siRNA was transfected with INTERFERin (Polyplus Transfection, France) according to the manufacturer’s recommendations. Total RNAs were extracted using the MACHEREY-NAGEL RNA extraction kit (REF 744352.4). A total of 500 ng of RNA was converted to first-strand cDNA using the RevertAid First Strand cDNA Synthesis Kit (Thermo Scientific, MA, USA). Real-time qPCRs were performed in 96-well plates using the IQ SYBR Green Supermix (Catalog #1708880, BioRad, CA, USA). Data were quantified using ΔΔ-Ct method and were normalized to GAPDH expression. siRNA and primers sequence used in the study are summarized (Table S4).

### HS27A cells differentiation, Alizarin Red staining and RNAseq

#### RNA samples

HS27A cells differentiation protocol toward osteoblasts was adapted from the team of Dr. Véronique Maguer-Satta (Cancer Research Center of Lyon -CRCL). HS27A cells were cultured in human Osteoblast differentiation media (417D-250) for 15 days. Cells undergo Alizarin red stain which labels calcium deposit to verify the differentiation. For staining, cells were rinsed in 1X PBS and fixed with 4% PFA for 30min. This is followed by washing cells with water and staining with 2% Alizarin solution (pH 4.2) for 2-3 min. RNA was extracted from cells at proliferating and differentiating states using MACHEREY-NAGEL RNA extraction kit (Ref 744352.4).

#### Construction of the libraries and sequencing

Determination of RNA concentration by fluorometric assay (Qubit4.0, Thermofisher) and analysis of distribution profiles of total RNA samples by capillary electrophoresis (Tapestation 4150, Thermofisher) were conducted by the Sequencing platform (PSI, IGFL, Lyon). The quality and quantity of the triplicate samples were satisfactory for initiating library construction, with no degradation of samples observed and RIN values > 8.6. mRNA capture was performed using the “NEBNext® Poly(A) mRNA Magnetic Isolation Module” protocol (New England Biolabs, E7490L) with a starting material of 1µg of total RNA for each sample, the various steps of the protocol having been carried out according to the supplier’s recommendations. Libraries were constructed using the Lexogen protocol “CORALL Total RNA-Seq Library Prep Kit with UDI 12 nt Set B1” protocol (Clinisciences, 118.96) according to the supplier’s recommendations and from 10μl of previously purified mRNA per sample. 16 cycles of PCR amplification were performed to finalize the construction of indexed library. The libraries were quantified (Qubit4.0, Thermofisher) and qualified (Tapestation 4150, Thermofisher). Sequencing was performed on the Illumina NextSeq 500 platform using Mid Output reagents with the following parameters: PE 2x81bp and 6bp index i7. A total of 194,006,581 reads were generated (Table S3).

#### Bioinformatics analyses

Reads were mapped onto the human genome using a local pipeline derived from the one supported by Lexogen (https://www.lexogen.com/corall-data-analysis/). Lexogen has made their pipeline available on the Lexogen’s Kangooroo app webpage (https://kangooroo.com/home). In brief, UMI were first trimmed with the program umi_tools extract (version 1.1.2, [51]). Adapter sequences were then removed using the Cutadapt package (version 3.5, [52]) and reads quality control was performed using FastQC (version 0.11.8; https://www.bioinformatics.babraham.ac.uk/projects/fastqc/). Read mapping was achieved with STAR (Spliced Transcripts Alignment to a Reference, version 2.6.1a, [53]) using the Homo sapiens GRCh38 genome reference (Ensembl, release 109). UMI deduplication was conducted with the program umi_tools dedup (version 1.1.2, [51]) enabling the accurate bioinformatic identification of PCR duplicates. The mapped reads were then quantified by Mix² (version 1.4.0.1, [54]), allowing to generate a count table per sample. Differential gene expression analysis was performed with DESeq2 [55] using a custom R script available on the following GitLab repository: https://gitbio.ens-lyon.fr/igfl/merabet/HS27ADiff. A Docker image is also provided to reproduce the results of the analysis.

#### xCELLigence Assays

The xCELLigence system (ACEA Biosciences Inc., San Diego, CA, USA), which records cellular events in real time by measuring electrical impedance across microelectrodes integrated on the bottom of culture plates (E-plates), was utilized in proliferation experiments. First, HS27A cell culture media were added to each well of 16-well E-plates (ACEA Biosciences Inc., San Diego, CA, USA) to measure background impedance. Then, 10.10^3^ of HS27A cells were transfected with a mixture of control, 14-3-3e, TAZ, SERPINB4 siRNAs at 10 pmol/mL using INTERFERin (Polyplus Transfection, France) according to the manufacturer’s protocol. Transfected cells were directly seeded on E-plates (5.10^5^ cells/well), and the impedance was measured every 15 min for 96 h. The impedance signal was proportional to the number of cells proliferating in each well and was displayed as cell index.

Data were analyzed with the RTCA Software 2.0 and were presented as mean +/− SEM of three experiments performed in triplicate.

---

**Supplementary Figure 1.**
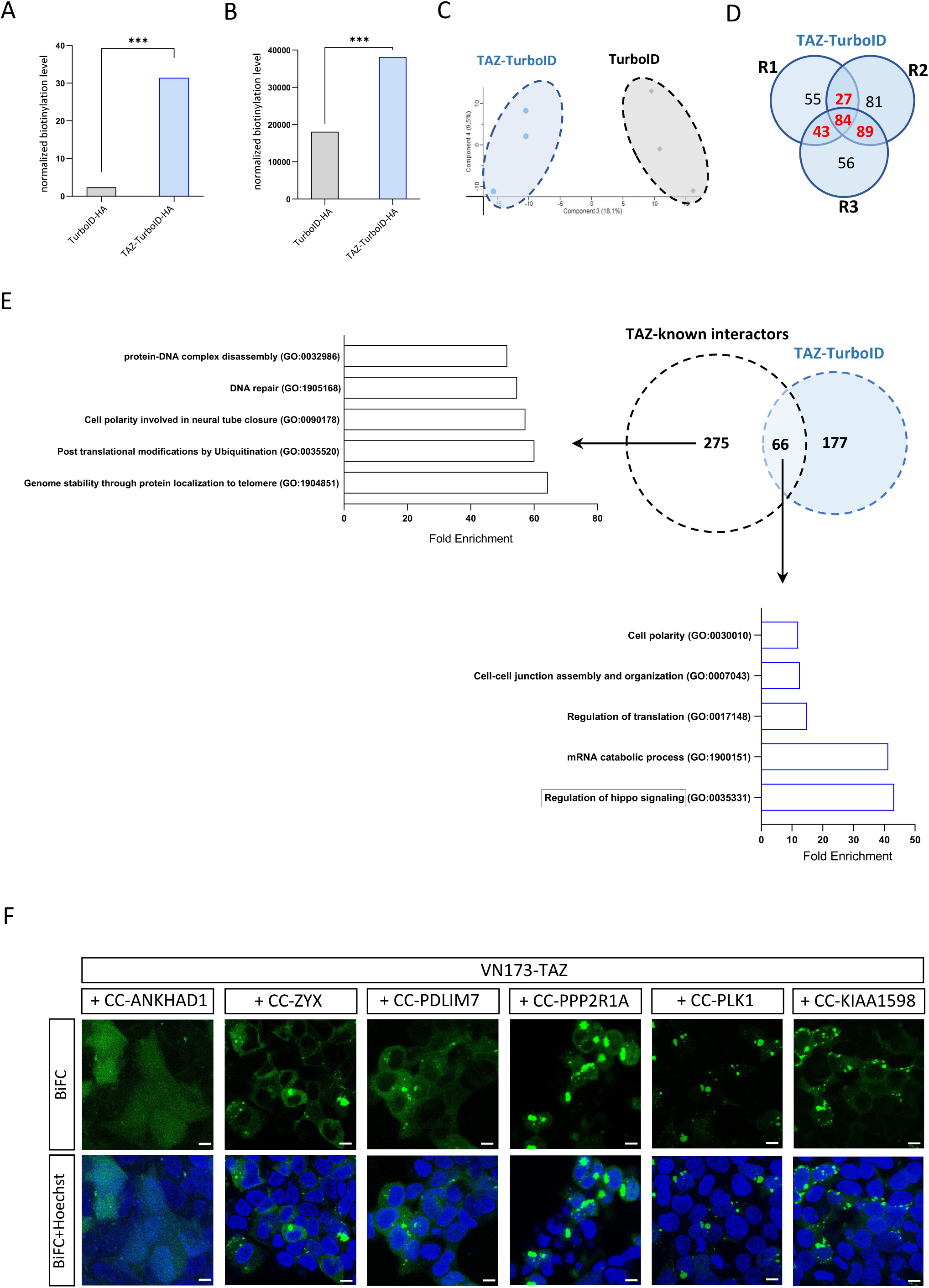
Biotinylation activity and interactome properties of TAZ-TurboID in HEK293T cells. **(A)** Quantification of the SA-555 immunostaining in HEK293T cells expressing TurboID or TAZ-TurboID, as indicated. Quantification was normalized over HA-staining. **(B)** Quantification of the intensity of bands revealed with the streptavidin staining for TurboID and TAZ-TurboID, as indicated. Quantification was normalized over HA-staining from the input. **(C)** Principal component analysis of TurboID and TAZ-TurboID replicates. **(D)** Venn diagram for TAZ-TurboID specific interactions (after filtering against TurboID) captured with the three biological replicates (R1-R3). **(E)** Go term analysis of the 66 common interactions between TAZ-TurboID and the known interactions of TAZ in the literature, and of the remaining 275 known interactions not captured in this study. **(F)** Illustrative confocal acquisitions of BiFC between VN-TAZ and new candidate partners already known in the literature and fused to CC, as indicated. Hoechst stains for nuclei in live HEK293T cells (blue). Scale bar = 20 µm.

**Supplementary Figure 2.**
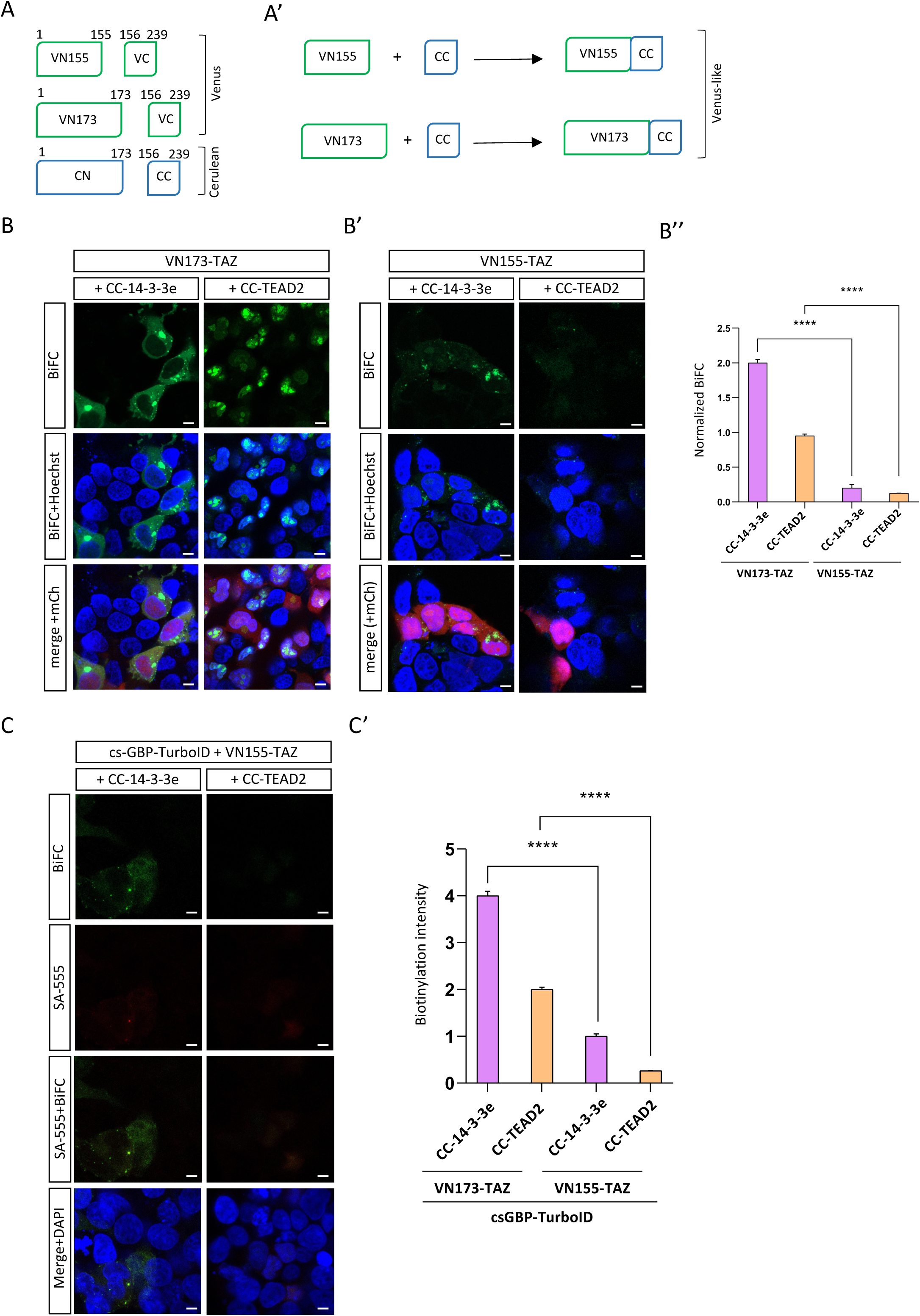
Comparison between VN173 and VN155 for doing Bi-nano-ID. **(A)** Scheme of the complementation between VN155 or VN173 from Venus and CC from Cerulean. The two complementation systems produce a Venus-like signal. **(B-B’)** Comparison of BiFC performed with VN173-TAZ (B) or VN155-TAZ (B’). BiFC was performed with CC-14-3-3e or CC-TEAD2 in live HEK293T cells. Hoechst stains for nuclei (blue) and mCherry (mCh) for transfection efficiency (red). **B’’** Quantification of BiFC obtained with VN173-TAZ or VN155-TAZ. BiFC was normalized over transfection efficiency in each condition. Quantification was deduced from at least three biological replicates. Statistical tests were performed with one-way ANOVA (\*\*\*\**p* < 0.0001). **(C)** Validation of the weak biotinylation activity of csGBP-TurboID when doing BiFC with VN155-TAZ. Biotinylation is revealed with streptavidin (SA) coupled to the 555 dye (red). Interaction between the fusion TAZ and 14-3-3e or TEAD2 is revealed by BiFC (green). Dapi stains for nuclei (blue). Scale bar = 20 µm. **(C’)** Quantification of the biotinylation signal for each condition, as indicated. Quantification was obtained from three biological replicates. Statistical tests were performed with one-way ANOVA (\*\*\*\**p* < 0.0001).

**Supplementary Figure 3.**
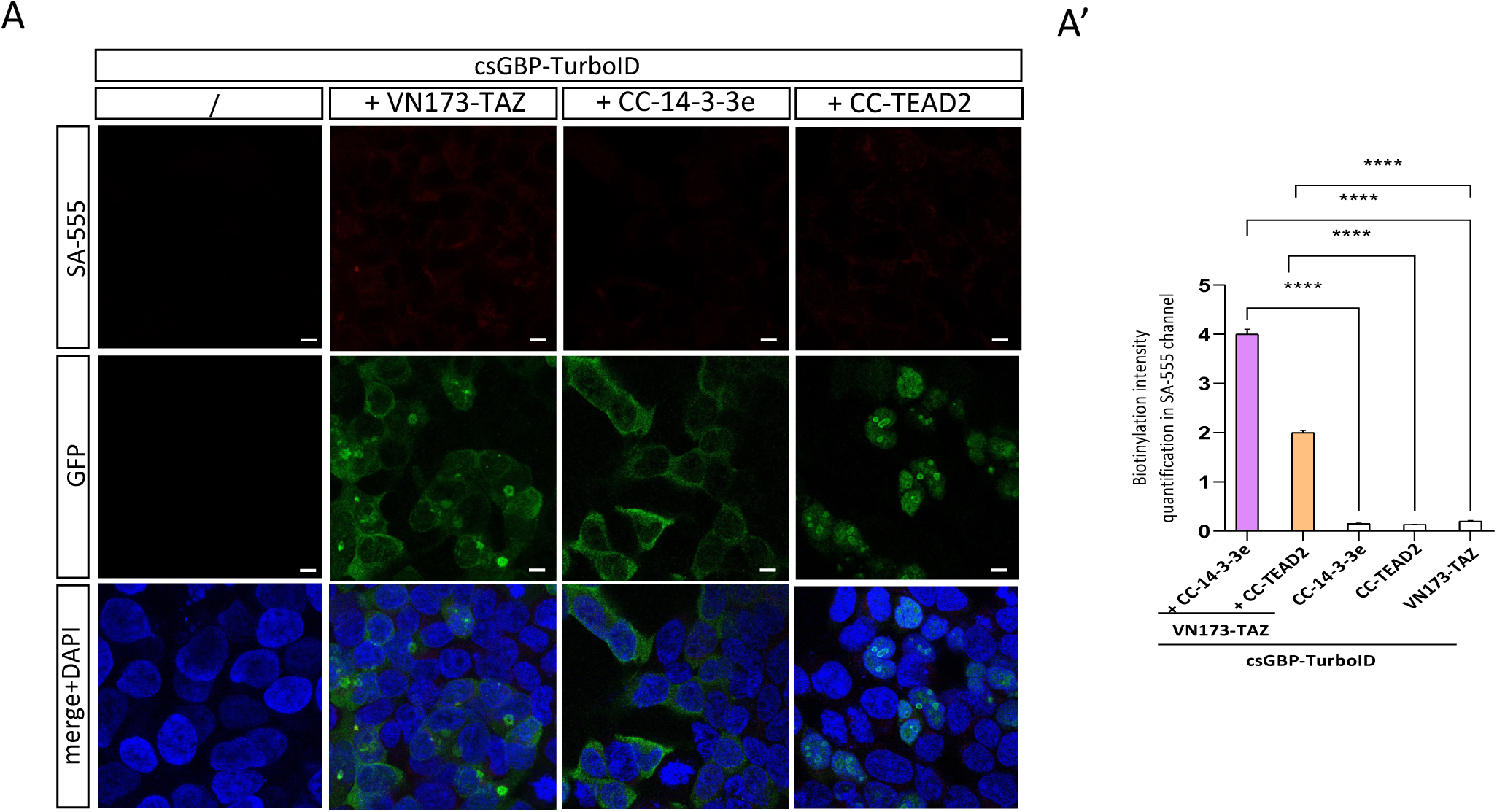
Specificity of the csGBP nanobody for the BiFC. **(A)** Fluorescent immunostaining with SA-555 (red) and anti-GFP (green) recognizing the VN and CC fragment with single construct transfection, as indicated. DAPI (blue) stains for nuclei. **(A’)** Quantification of the biotinylation signal when compared to signals obtained with BiFC complexes. Quantification was obtained from three biological replicates. Statistical tests were performed with one-way ANOVA (\*\*\*\**p* < 0.0001).

**Supplementary Figure 4.**
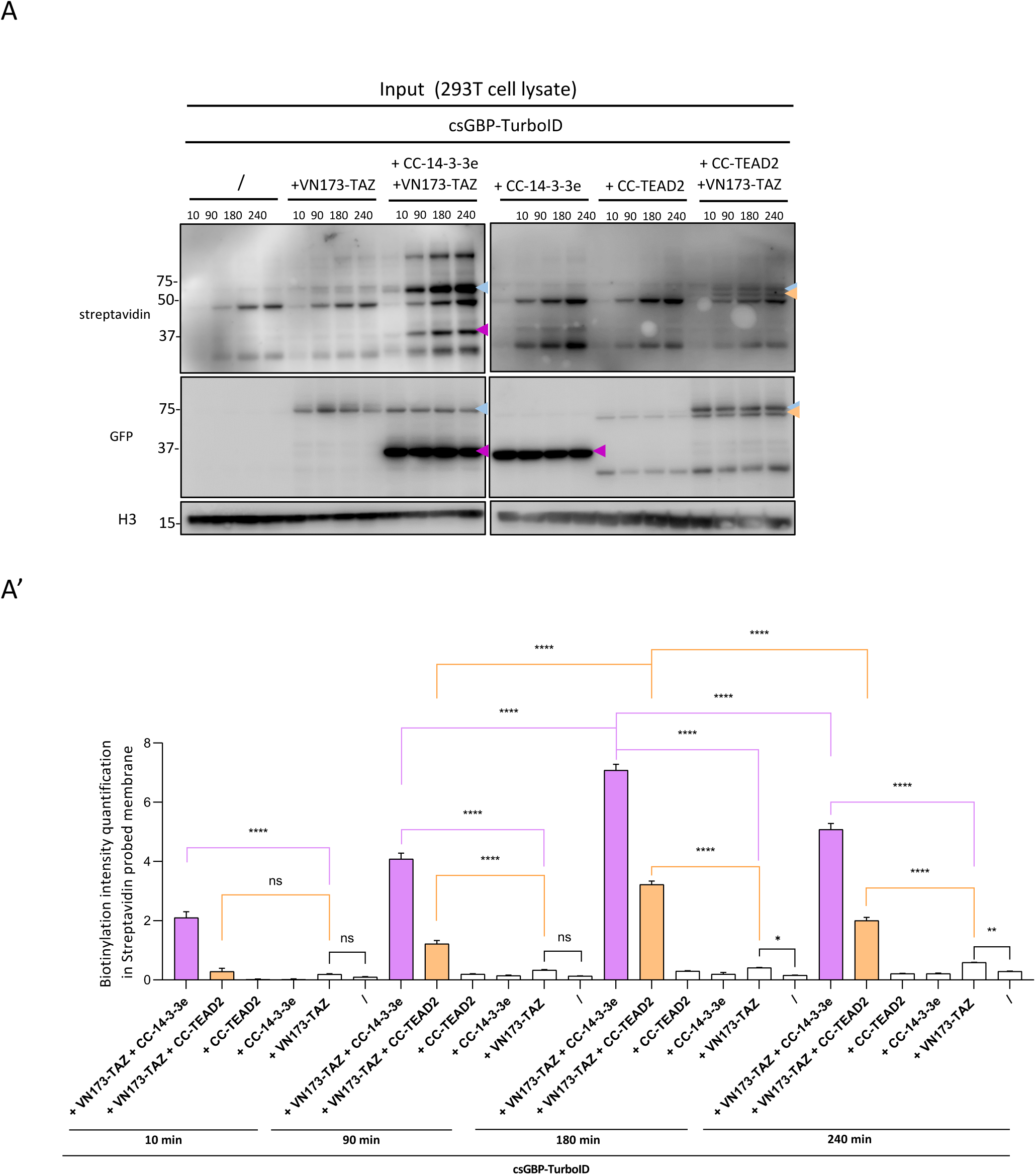
Western blots of biotinylated proteins extracted from HEK293T cells upon transfection with the different constructs, as indicated. **(A)** Input fraction of extracted proteins revealed on the gel probed by streptavidin, GFP and H3 antibodies. Colored arrowheads indicate the band corresponding to VN173-TAZ (blue arrowhead), CC-14-3-3e (magenta arrowhead) and CC-TEAD2 (orange arrowhead). **(A’)** Histogram showing the quantification of the biotinylation bands in each condition, as indicated. The quantification was deduced by subtracting the AP band with the corresponding input band. It also took into account the non-specific AP bands of the nanobody for quantifying biotinylation of VN-TAZ and CC-TEAD2 (see Fig. 2). Statistical tests were performed with one-way ANOVA from three biological replicates (\*\*\*\**p* < 0.0001, \**p* < 0.05).

**Supplementary Figure 5.**
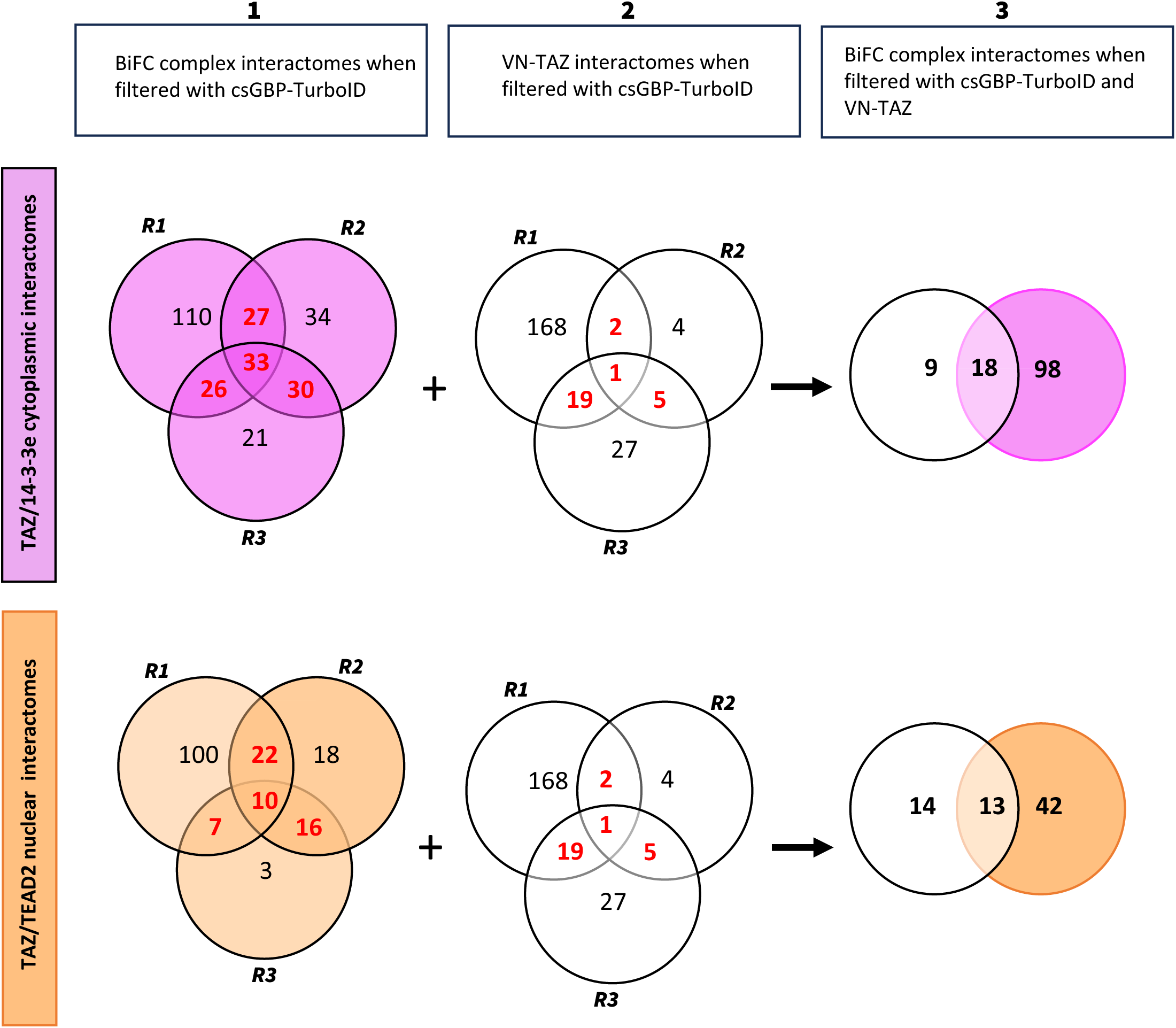
Filtering steps for identifying TAZ/14-3-3e and TAZ/TEAD2 interactomes. Filter 1 was applied to remove BiFC interaction candidates that were also found with dsGBP-TurboID alone (comparing BiFC+csGBP-TurboID to csGBP-TurboID condition in each replicate). Filter 2 identified interactions obtained with VN173-TAZ alone when compared to the dsGBP-TurboID condition in each replicate. Filter 3 identified BiFC-specific interactions when removing interactions found with csGBP-TurboID alone (filter 1) and VN-TAZ alone (filter 2). Only interactions found in 2 or 3 replicates were considered. This filtering was applied for TAZ/14-3-3e (upper panel) and TAZ/TEAD2 (lower panel) complexes, leading to the identification of 98 and 42 interactions, respectively.

**Supplementary Figure 6.**
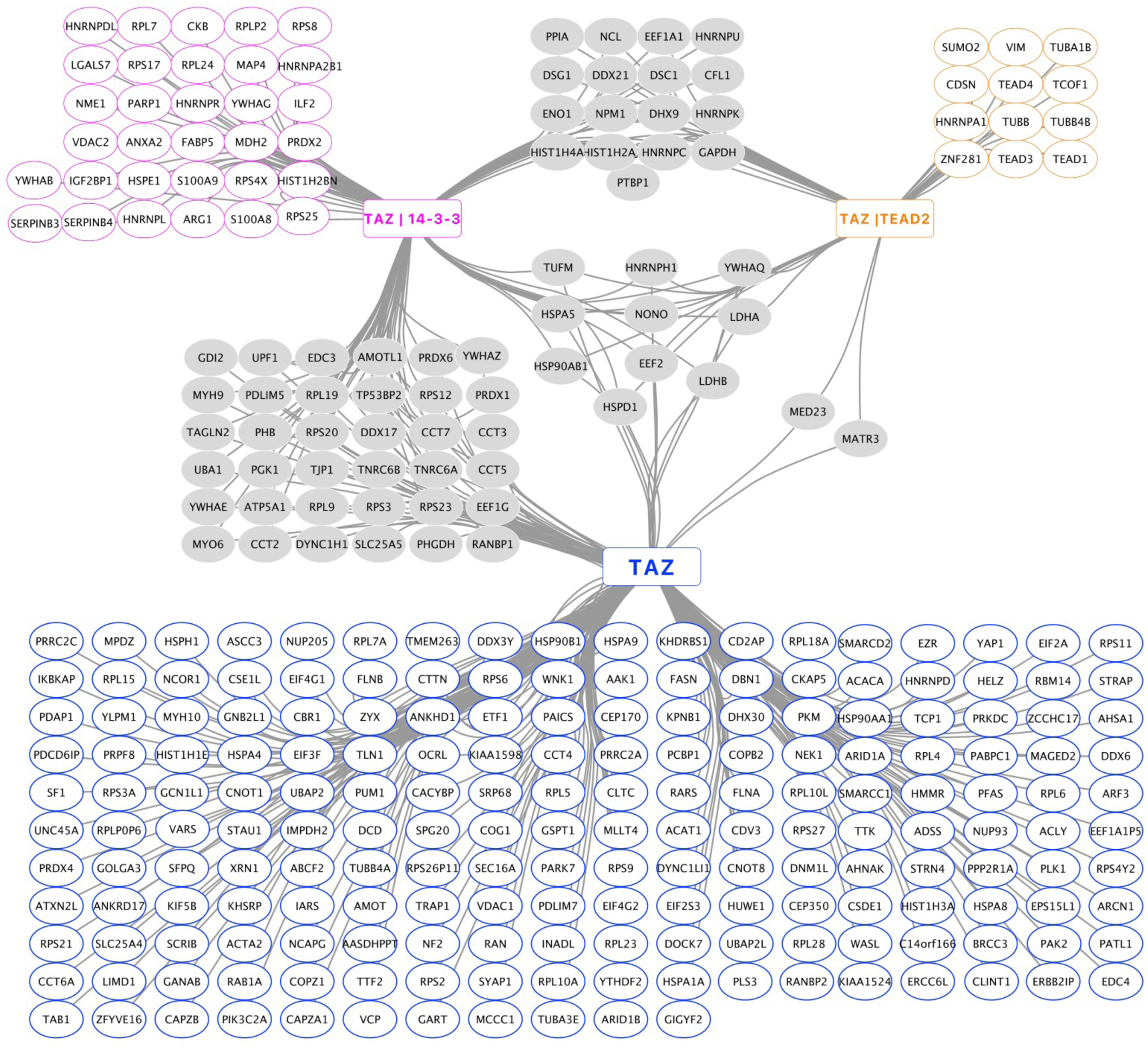
Molecular clustering of TAZ-TurboID, TAZ/14-3-3e and TAZ/TEAD2 interactomes.

**Supplementary Figure 7.**
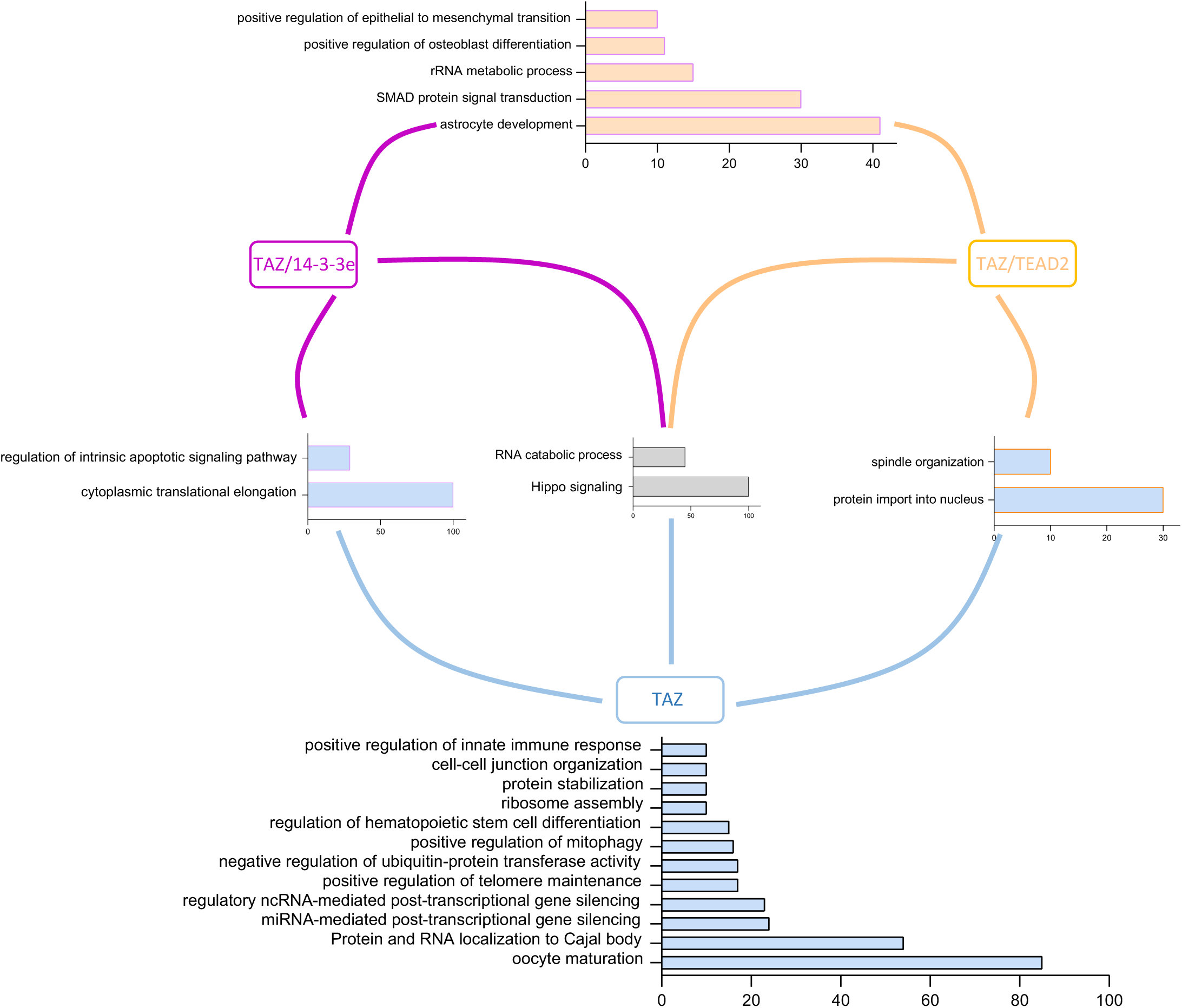
Functional clustering of TAZ-TurboID, TAZ/14-3-3e and TAZ/TEAD2 interactomes. The clustering was based on GO terms that were 10 times enriched at the minimum. Specific TAZ/14-3-3e and TAZ/TEAD2 Go term enrichments are shown in the Fig. 3.

**Supplementary Figure 8.**
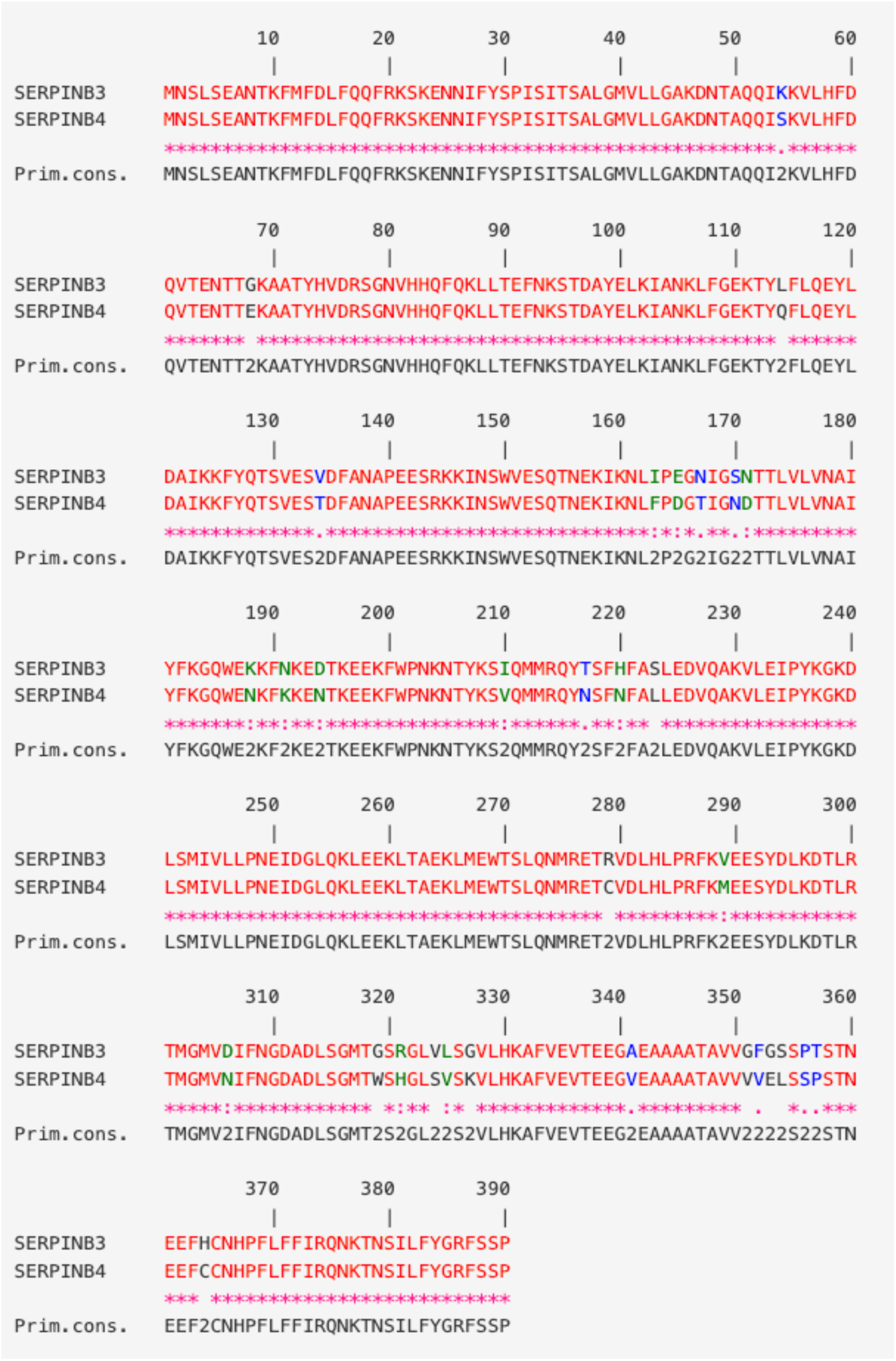
Alignment of SERPINB3 and SERPINB4 protein sequences. Alignment was obtained by using the CLUSTALW web tool from PRABI (https://prabi.ibcp.fr/htm/site/web/app.php/home).

**Supplementary Figure 9.**
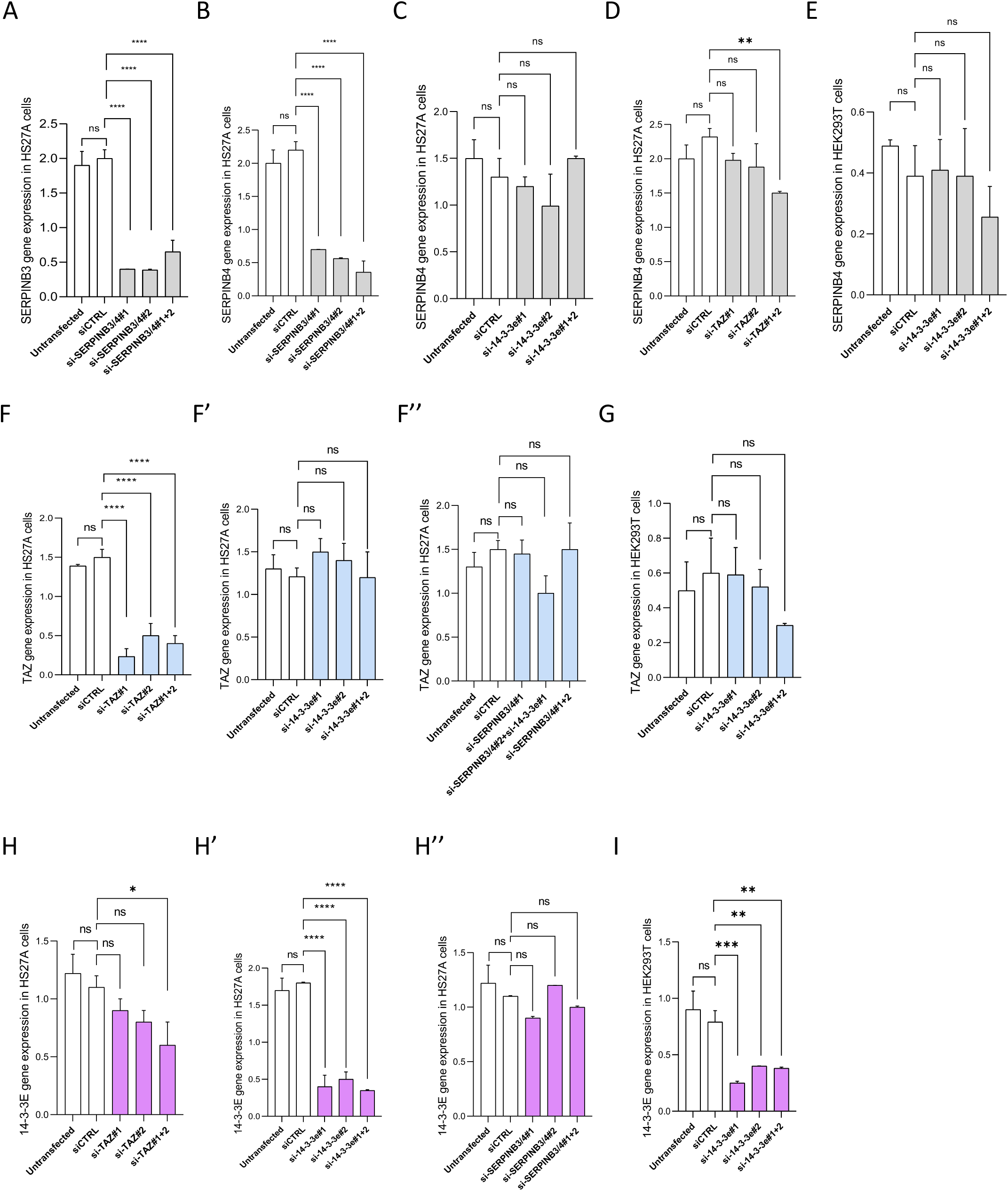
Effects of siRNAs. **(A)** Effect of siRNAs targeting *SERPINB3* and *SERPINB4* on endogenous *SERPINB3* in HS27A cells. **(B)** Effect of siRNAs targeting *SERPINB3* and *SERPINB4* on endogenous *SERPINB4* in HS27A cells. **(C)** Effect of siRNAs targeting *14-3-3* on endogenous *SERPINB4* expression in HS27A cells. **(D)** Effect of siRNAs targeting *TAZ* on endogenous *SERPINB4* expression in HS27A cells. **(E)** Effect of siRNAs targeting *14-3-3* on endogenous *SERPINB4* expression in HEK293T cells. **(F-F’’)** Effects of siRNAs targeting *TAZ* (F), *14-3-3e* (F’) or *SERPINB3/B4* (F’’) on endogenous *TAZ* expression in HS27A cells. **(G)** Effects of siRNAs targeting *14-3-3e* on endogenous *TAZ* expression in HEK293T cells. **(H-H’’)** Effects of siRNAs targeting *TAZ* (H), *14-3-3e* (H’) or *SERPINB3/B4* (H’’) on endogenous *14-3-3e* expression in HS27A cells. **(I)** Effects of siRNAs targeting *14-3-3e* on endogenous *14-3-3e* expression in HEK293T cells. Graphs result from the quantification of RTqPCR from three biological replicates in each condition. Statistical tests were performed with one-way ANOVA (ns: non-significant, \*\**p* < 0.01, \*\*\*\**p* < 0.0001). Note that *SERPINB3* and *SERPINB4* are 98% identical at the nucleotide sequence level.

**Supplementary Figure 10.**
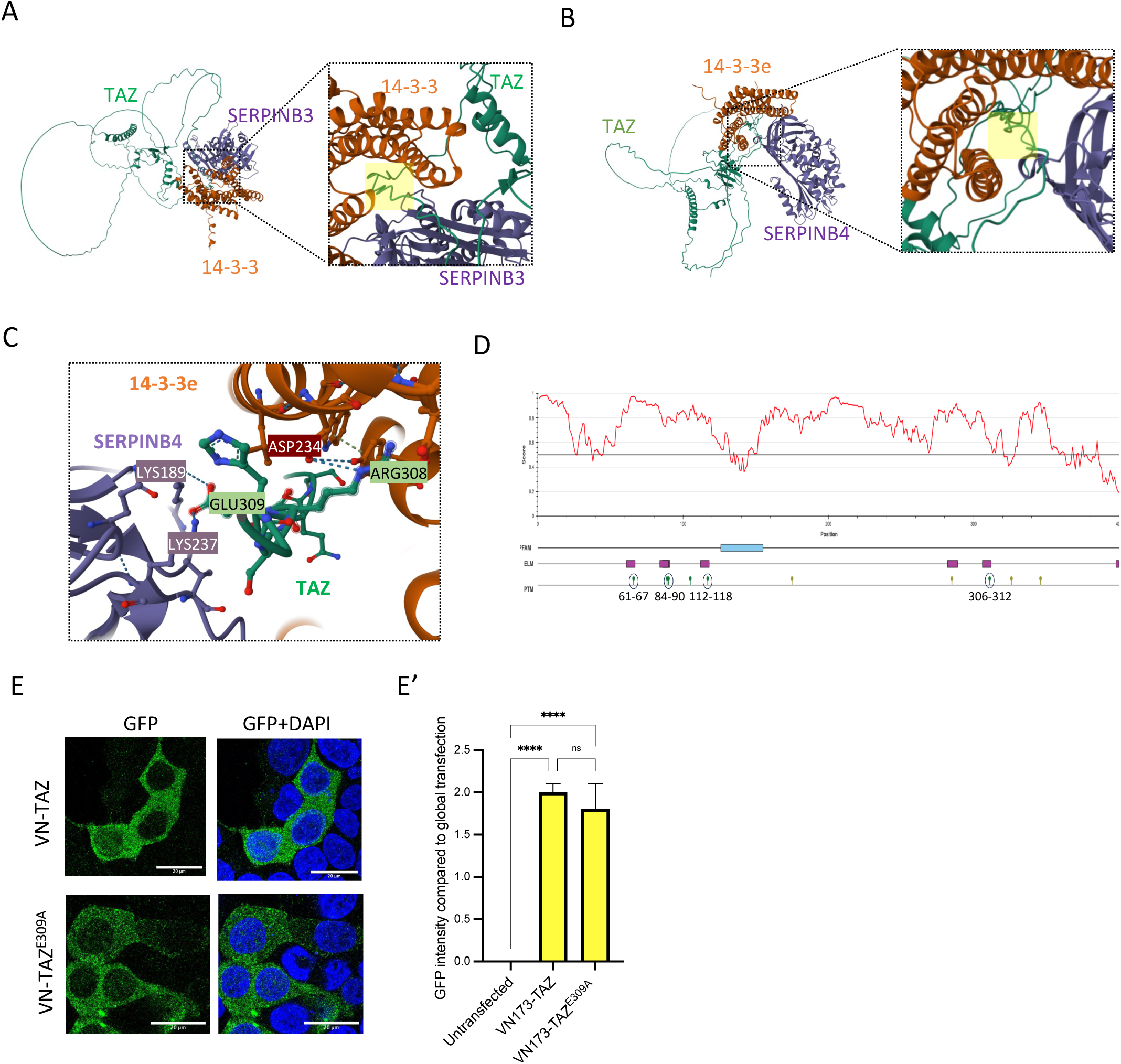
Structure and interaction predictions for TAZ, 14-3-3e, SERPINB3 and SERPINB4. **(A)** AlphaFold multimer prediction of TAZ/14-3-3e/SERPINB3 trimeric protein complex. **(B)** AlphaFold multimer prediction of TAZ/14-3-3e/SERPINB4 trimeric protein complex. Enlargements highlight the disordered C-terminal region of TAZ in contact with SERPINB3 (A) or SERPINB4 (B) and 14-3-3e (highlighted with a light-yellow box). **(C)** Enlargement at the level of the Glu309 residue of TAZ that makes contacts with two Lys residues of SERPINB4. Among other residues in contact with 14-3-3e in the same region, the residue Arg308 of TAZ interacts with the residue Asp234 of 14-3-3e. **(D)** Prediction of disordered regions (red line above the gray line), ordered domain (blue box), short linear interaction motifs (violet boxes) and LATS1-phosphorylation sites in TAZ protein sequence. Adapted from ELM (http://elm.eu.org/search.html) and IUPred2A (https://iupred2a.elte.hu/). **(E-E’)** Quantification of wild type or mutated TAZ expression in HEK293T cells. VN-fusion constructs were revealed with anti-GFP (green). Dapi stains for nuclei. Statistical tests were performed with one-way ANOVA (\*\*\*\**p* < 0.0001).

**Supplementary Figure 11.**
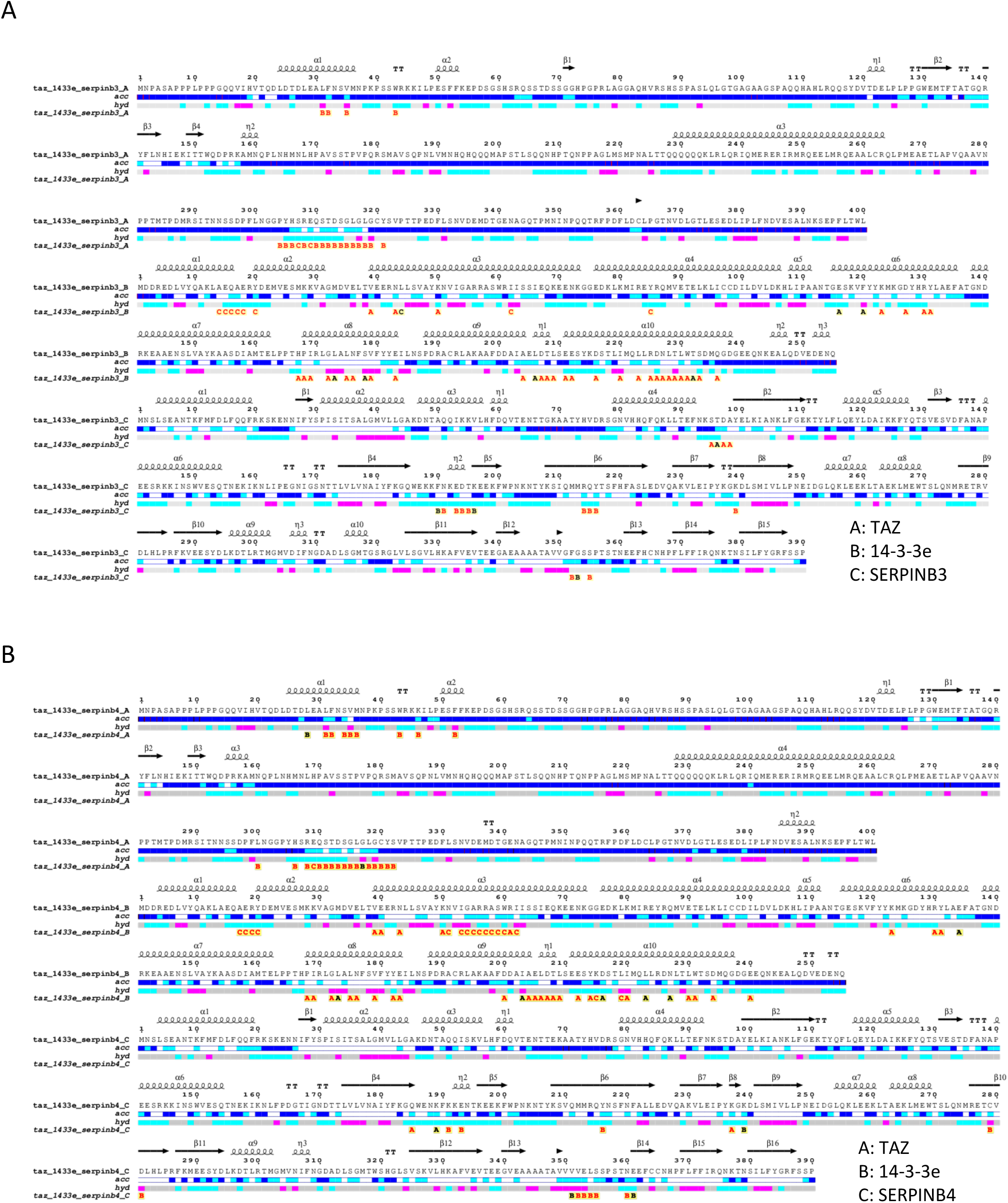
Endscript2 analyses of interaction contacts predicted in the TAZ/14-3-3-3e/SERPINB3 (A) and TAZ/14-3-3e/SERPINB4 (B) complexes. The sequences of TAZ (Chain A) 14-3-3e (chain B) and SERPINB3/4 (chain C) are presented from top to bottom. They are adorned with secondary structure elements, alpha-helices being represented by squiggles and beta-strands by arrows. Accessibility bar (acc, blue accessible, cyan intermediate, white buried) hydropathy bar (hyd, cyan hydrophylic, pink hydrophobic) and intermolecular contact letters (A to C refers to the chain of the residue contacted; a red letter indicates a contact distance < 3.2 Å and a black letter a contact distance in the range 3.2-3.7 Å) are written below the sequences.

**Supplementary Figure 12.**
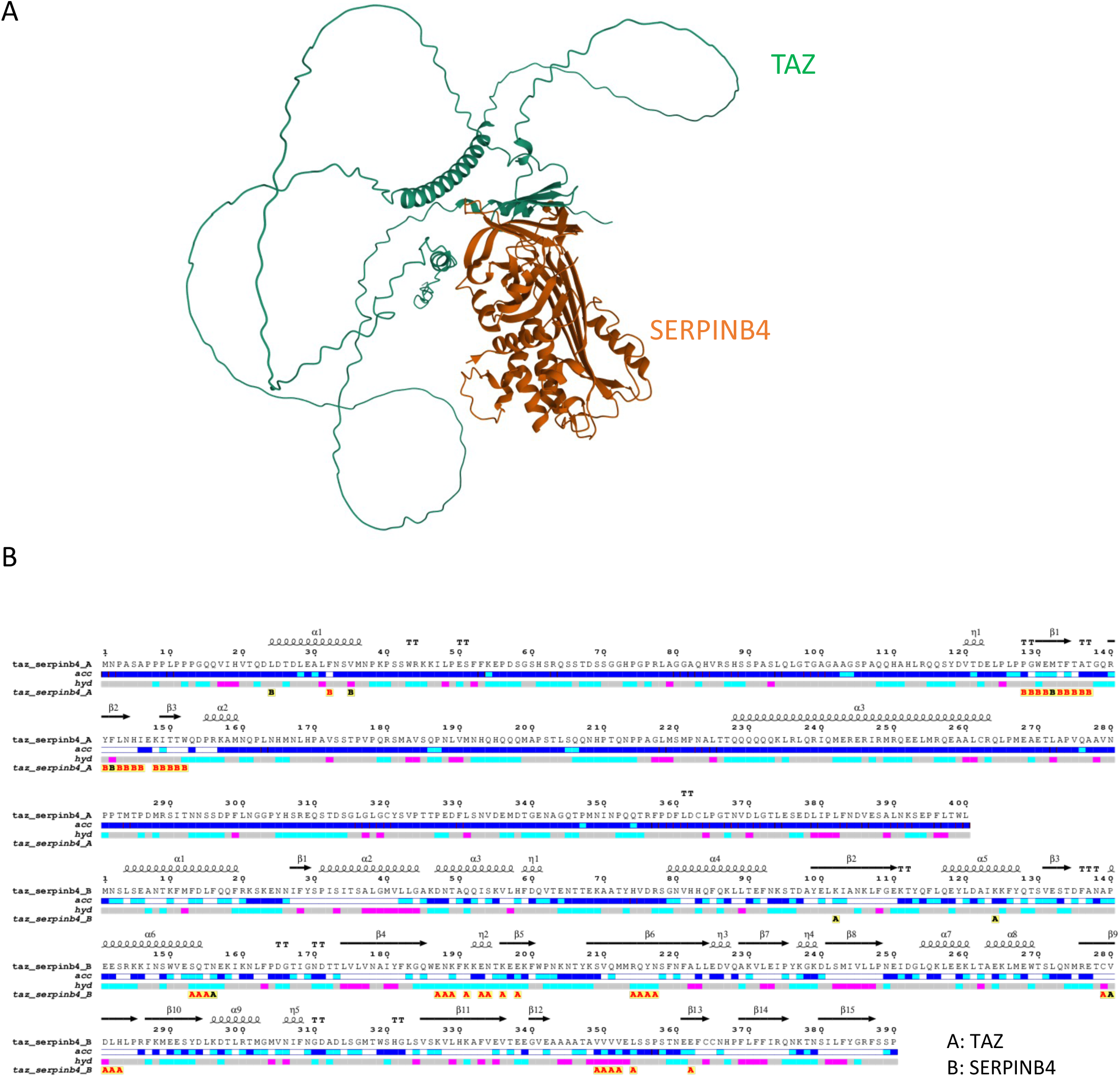
AlphaFold prediction (A) and Endscript2 annotations (B) of TAZ/SERPINB4 dimeric protein complex.

**Supplementary Figure 13.**
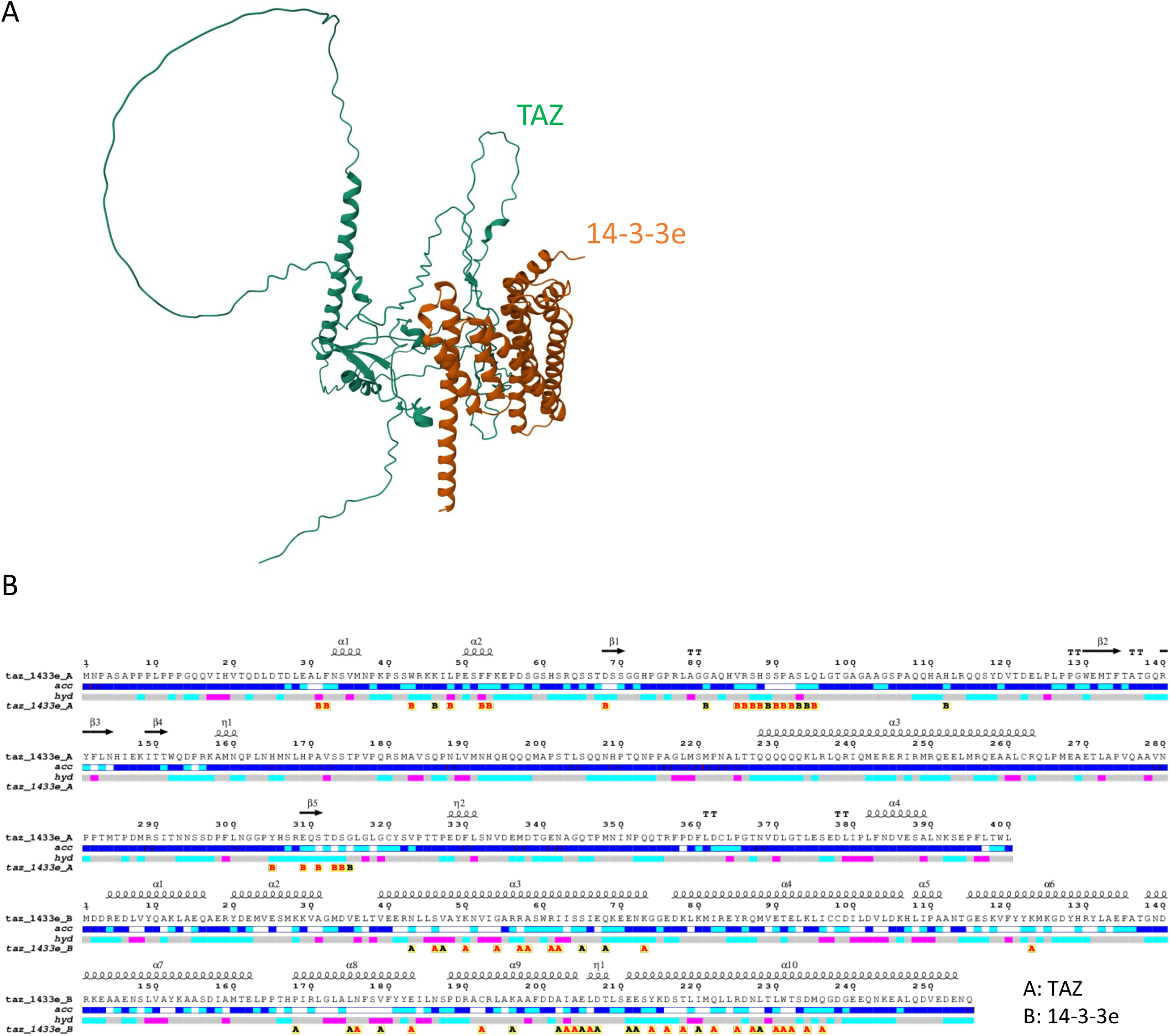
AlphaFold prediction (A) and Endscript2 annotations (B) of TAZ/14-3-3e dimeric protein complex.

**Supplementary Figure 14.**
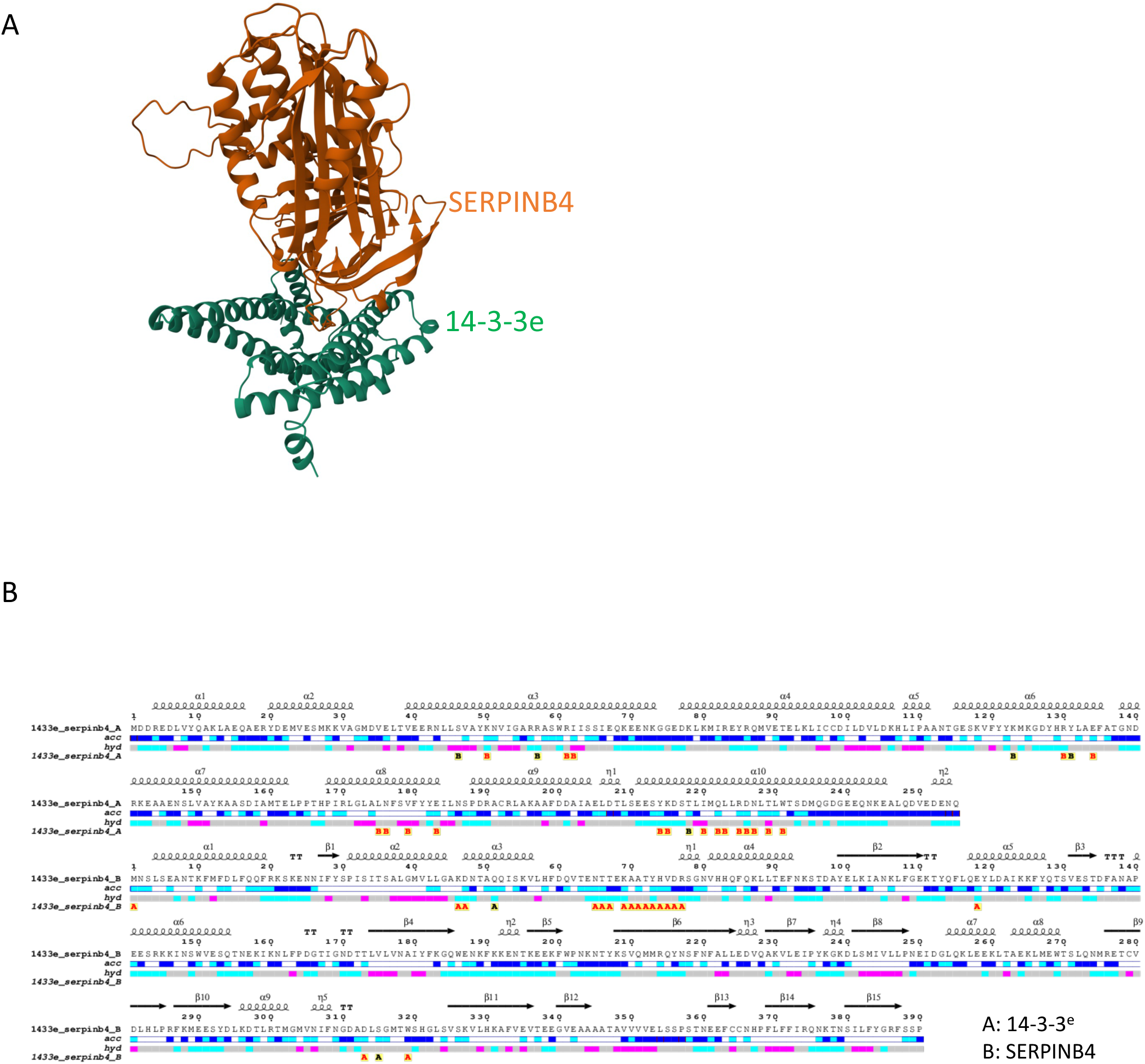
AlphaFold prediction (A) and Endscript2 annotations (B) of 14-3-3e/SERPINB4 dimeric protein complex.

**Supplementary Tables Supplementary Table 1. Mass spectrometry data and GO term enrichment for TAZ-TurboID**

**Supplementary Table 2. Mass spectrometry data and GO term enrichment for TAZ/14-3-3e and TAZ/TEAD2 complexes**

**Supplementary Table 3. RNA-seq data of HS27A cells.**

**Supplementary Table 4. Sequences of oligonucleotides and primers used in this study for cloning and RTqPCRs**

## Authors contribution

S.M. designed the study. N.H.S. performed the experiments and acquired the data. J.C. and N.H.S., with the support of S.M. and F.D., were responsible of proteomics data analyses. N.H.S and L.G. analyzed the RNA-seq data. Y.J. was responsible for the cytoscape network illustration. S.M. and N.H.S. wrote the paper and all authors amended it.

## Acknowledgments

We are beyond grateful for the help of Christelle Forcet in performing the proliferation assay and support at the experimental level. We thank also Benjamin Gillet, Sandrine Hughes and Julien Dellinger for the RNA-seq preparation and valuable help in the transcriptomic analyses. We thank Adeline PAGE from the Protein Science Facility at the SFR bioscience (UAR3444/CNRS, US8/Inserm, ENS de Lyon) for Mass spectrometry analyses. We are very grateful for the help provided by Sylvain Lefort and the team of Dr. Veronique Maguer-Satta (CRCL) for HS27A line and culturing tips. We thank Alain Guignandon and Daniel Bouvard for their valuable inputs in the project. We thank Julien Béthune for careful reading of the manuscript and helpful comments. This project was supported by Ambition Pack from the Region Rhone Alpes and the Foundation of Medical Research (FRM #FDT202304016377, 4th year grant for N.H.S).

